# Enhanced photosynthetic efficiency for increased carbon assimilation and woody biomass production in hybrid poplar INRA 717-1B4

**DOI:** 10.1101/2022.02.16.480797

**Authors:** Living Carbon Team, Yumin Tao, Li-Wei Chiu, Jacob W. Hoyle, Jessica Du, Karli Rasmussen, Patrick Mellor, Christian Richey, Julie Kuiper, Madeline Fried, Rebecca A. Dewhirst, Dominick Tucker, Alex Crites, Gary A. Orr, Matthew J. Heckert, Damaris G. Vidal, Martha L. Orosco-Cardenas, Madeline E. Hall

## Abstract

Increasing CO_2_ levels in the atmosphere and the resulting negative impacts on climate change have compelled global efforts to achieve carbon neutrality or negativity. Most such efforts focus on carbon sequestration through chemical or physical approaches. We aim to harness the power of synthetic biology to enhance plants’ natural ability to draw down and sequester carbon, thereby positively affecting climate change. Past decades of scientific progress have shed light on strategies to overcome the intrinsic limitations of carbon drawdown and fixation through photosynthesis, particularly in row crops in hopes of improving agricultural productivity for food security. Incorporating a photorespiration bypass in C3 plants has shown promising results of increased biomass and grain yield. Despite their globally dominant role in atmospheric carbon flux, the drawdown rates of most trees are currently limited by their C3 photosynthetic metabolism, and efforts to improve the photosynthetic capacity of trees, such as by reducing energy loss in photorespiration, are currently lacking. Here, we selected a photorespiration bypass pathway and tested its effectiveness on photosynthetic enhancement in hybrid poplar INRA717-IB4. The design includes a RNAi strategy to reduce the transportation of the photorespiration byproduct, glycolate, out of chloroplast and a shunt pathway to metabolize the retained glycolate back to CO_2_ for fixation through the Calvin-Benson cycle. Molecular and physiological data collected from two repeated growth experiments indicates that transgenic plants expressing genes in the photorespiration bypass pathway have increased photosynthetic efficiency, leading to faster plant growth and elevated biomass production. One lead transgenic event accumulated 53% more above-ground dry biomass over a five month growth period in a controlled environment. Pilot projects with photosynthesis-enhanced trees in the field are in progress. Our results provide a proof-of-concept for engineering trees to help combat climate change.

## INTRODUCTION

Originating from single-celled prokaryotes in the ocean approximately 2.5 billion years ago, photosynthesis is a natural process in plants consisting of a series of well-coordinated biochemical reactions that use sunlight energy to reduce atmospheric CO_2_ into energy-rich carbohydrates, while releasing oxygen back into the atmosphere. Both compounds are vital to most living organisms, including humans. Plants evolved to center the activities of photosynthesis in organelles known as chloroplasts. In chloroplasts sunlight is captured by the pigment chlorophyll, which energizes electrons derived from water molecules, while releasing oxygen in the thylakoid membrane. These high-energy electrons are then transferred through a series of carriers in photosystem I, photosystem II, cytochrome *bf* complex, and eventually drives the synthesis of ATP and NADPH by ATP synthase and NADP reductase located in the thylakoid membrane. In the stroma of a chloroplast, CO_2_ is fixed into carbohydrates *via* the Calvin-Benson cycle utilizing the ATP and NADPH produced during the aforementioned light reactions in the thylakoid membrane. The first step of the Calvin-Benson cycle is catalyzed by the enzyme RuBisCo, **r**ib**u**lose-1,5-**bis**phosphate **c**arboxylase/**o**xygenase, which as the name suggests can accept either CO_2_ or O_2_ as substrates to carry out either a carboxylation or oxygenation reaction, respectively. The carboxylation reaction leads to carbon fixation while oxygenation reaction leads to photorespiration. The fixed CO_2_ molecule moves on to anabolic pathways for sucrose and starch biosynthesis. The oxygenation product phosphoglycolate must be converted in a series of reactions that eventually generates one molecule of 3-phosphoglycerate and one molecule of CO_2_ (for more details on photosynthesis, see Johnson, 2016).

Numerous attempts utilizing a multitude of approaches, such as computational metabolic modeling, have been made to improve upon the inefficiencies of photosynthesis due to photorespiration (Xin et al., 2015). Recent advancement in synthetic biology techniques has contributed to this field of research through enzyme engineering and alternative pathway engineering (Kubis and Bar-Even, 2019). Photorespiration starts with the oxygenation activity of RuBisCo, especially when CO_2_ concentration is low. Plants grown at elevated CO_2_ concentrations have been shown to increase productivity (Ainsworth, 2008). Such results boosted the belief that engineering the RuBisCo enzyme on substrate specificity towards CO_2_ could substantially improve photosynthetic efficiency leading to higher productivity (Dhingra et al., 2004; Spreitzer et al., 2005; Whitney and Sharwood, 2007). However, enzyme engineering approaches to improve Rubisco catalysis by random or site-directed mutagenesis or directed evolution have largely failed to yield substantial kinetic enhancements (Spreitzer et al., 2005; Mueller-Cajar and Whitney, 2008; Whitney et al., 2011; Wilson et al., 2016). The lack of considerable improvements from these studies is potentially due to the existence of a trade-off between CO_2_ specificity and carboxylation velocity - such that carboxylation activity decreases when CO_2_ specificity is increased (Savir et al., 2010; Galmés et al., 2014). However, Young et al. (2016) evaluated the diversity of Rubisco kinetics in 11 different diatom algae species and found no statistically significant relationship between carboxylase efficiencies and CO_2_ specificities in diatom RuBisCo enzymes. Computational modeling analysis of RuBisCo kinetics also challenged the trade-off idea (Cummins et al., 2018). It remains a possibility that RuBisCo can be improved *via* enzyme engineering with a better designed screening system. Interestingly, Long et al. (2018) reported their attempt to assemble a simplified α-carboxysome in tobacco chloroplasts by replacing native Rubisco with the large and small subunits of Rubisco from cyanobacteria, along with two key structural subunits with better carboxylase efficiency and substrate specificity. Successful expression of the functional α-carboxysome in tobacco chloroplasts led to an increase in biomass production. Other attempts have been on improving enzymes in the Calvin-Benson cycle to enhance the rate of RuBP regeneration which is known to limit the carbon fixation rate under certain conditions. Computational models have suggested that the natural distribution of enzymes within the Calvin-Benson cycle are suboptimal (Zhu et al., 2007), especially under elevated CO_2_ concentrations. Overexpression of sedoheptulose 1,7-bisphosphatase or fructose-1,6-bisphosphate aldolase in N. tabacum resulted in a higher carbon fixation rate and increased biomass at elevated CO_2_ levels (Rosenthal et al., 2011; Uematsu et al., 2012). Even at ambient CO_2_ concentration, overexpression of limiting enzymes in the Calvin cycle was shown to boost carbon fixation in tobacco (Miyagawa et al., 2001; Lefebvre et al., 2005; Simkin et al., 2015).

The next obvious approach centered around engineering photorespiration by-pass pathways to metabolize the byproduct of RuBisCo oxygenation, phosphoglycolate, preventing it from entering the natural photorespiratory pathway thus minimizing the loss of CO_2_ and energy. The first such bypass pathway was to engineer into Arabidopsis an *E. coli* glycolate catabolic pathway consisting of three enzymes: glycolate dehydrogenase, glyoxylate carboligase, and tartronic semialdehyde reductase (Kebeish et al., 2007). The design was to mimic the natural photorespiration process of returning 75% glycolate to the Calvin-Benson cycle without ammonia production. Increases in photosynthesis rate and biomass were observed in transgenic Arabidopsis. Carvalho (2011) designed a bypass to convert glycolate into hydroxypyruvate in peroxisomes by two enzymes from *E. coli:* glyoxylate carboligase and hydroxypyruvate isomerase. This bypass releases CO_2_ in the peroxisome and does not produce extra ammonia. However, transgenic tobacco showed stunted growth in ambient air (Carvalho, 2011). Maier et al. (2012) reported another bypass design to completely convert glycolate into CO_2_ in the chloroplast. The pathway consisted of three new enzymes: the Arabidopsis glycolate oxidase (directing the transgenic enzyme into chloroplast instead of peroxisome location for the endogenous one), pumpkin malate synthase, and the *E. coli* catalase. Together with two endogenous enzymes (NADP-malic enzyme and pyruvate dehydrogenase), this pathway mimics the *E. coli* glycolate oxidative cycle in Arabidopsis chloroplasts. Increased photosynthesis rate and dry weight biomass were observed (Maier et al. 2012). Nolke et al. (2014) expressed a polyprotein comprising all three subunits of E. coli glycolate dehydrogenase in plastids of transgenic potato. Transgenic lines with the highest polyprotein levels and the glycolate dehydrogenase activity produced significantly higher levels of carbohydrate, resulting in a substantial increase in shoot and leaf biomass and tuber yield. Recently, South et al. (2019) reported yet another bypass design that not only has a shunt pathway to metabolize glycolate utilizing the glycolate dehydrogenase from *Chlamydomonas reinhardtii* and malate synthase from *Cucurbita maxima*, but also includes a RNAi design to inhibit the plastidial glycolate transporter, Plgg1, in an attempt to limit the entrance of glycolate into photorespiration pathway. With a successful reduction in the expression of tobacco Plgg1, transgenic plants showed biomass increase as high as 40% in field conditions (South et al. 2019). The effectiveness of photorespiration by-pass pathway designs to enhance photosynthesis efficiency has recently been tested in one crop species, rice. The first design, called GOC bypass, consists of three rice enzymes: **g**lycolate oxidase, **o**xalate oxidase, and **c**atalase (Shen et al. 2019). Similar to the idea of Maier et al. (2012) to completely metabolize glycolate into CO_2_ in the chloroplast, this bypass design uses three rice enzymes. Increases in photosynthetic rate and biomass were observed in transgenic rice, especially under high light conditions, but seed setting was reduced (Shen et al., 2019). Wang et al. (2020) reported a GCGT bypass design with four genes: a rice glycolate oxidase and three *E. coli* enzymes including the catalase, the glyoxylate carboligase, and the tartronic semialdehyde reductase. This bypass expects greater increase in photosynthetic rate and grain yield by mimicking the natural 75% returning of glycolate to Calvin cycle as opposed to 100% conversion into CO_2_. Higher increases in photosynthesis rate, biomass, and grain yield were observed. However, like the GOC bypass, seed setting is reduced in transgenic rice.

Other attempts harness the power of enzyme and metabolic engineering to develop synthetic routes that bypass photorespiration without CO_2_ release. Trudeau et al. (2018) used computational design and directed evolution to establish this activity in two sequential reactions. An acetyl-CoA synthetase was engineered for higher stability and glycolyl-CoA synthesis. A subsequent propionyl-CoA reductase was engineered for higher affinity of glycolyl-CoA and selectivity of NADPH over NAD+, thereby favoring reduction over oxidation. The combination of these evolved enzymes with pre existing ones supported the *in vitro* recycling of glycolate to RuBP without the loss of CO_2_, providing proof-of-concept for carbon-conserving photorespiration (Trudeau et al., 2018). Yu et al. (2018) designed the MCG pathway that can produce acetyl-CoA from C3 sugars without releasing CO_2_ which, when coupled with glycolate dehydrogenase, can assimilate glycolate to acetyl-CoA without net carbon loss. López-Calcagno et al. (2019) reported that leaf specific overexpression of the H-protein of the glycine cleavage system increased biomass yield in transgenic tobacco plants over two seasons of field growth.

In trees, however, there have rarely been any attempts to increase woody biomass *via* manipulations of enzymes or metabolic pathways for photosynthesis enhancement. Living Carbon strives to create the next generation of trees *via* synthetic biology approaches with the goal of increasing carbon assimilation and sequestration in an attempt to help combat the looming consequences of climate change. We take a holistic view of overall carbon capture and sequestration, employing multidisciplinary approaches to work on areas including photosynthesis enhancement, growth rate increase, heavy metal accumulation, fungal disease resistance, drought tolerance, salt tolerance, phytoremediation, and prolonged and permanent carbon storage. Built upon the tremendous amount of knowledge from decades of research progress made by the greater scientific community, and through the combined efforts of increasing carbon drawdown from the atmosphere into plants and decreasing the release of CO_2_ back into the air, we hope to greatly aid the world to achieve its goal of carbon neutrality or carbon negativity *via* living organisms. This paper summarizes our work on one specific approach which is to engineer a photorespiration bypass to enhance carbon fixation in hybrid poplar trees.

## MATERIAL & METHODS

### Poplar plant growth and propagation

Hybrid *Populus tremula* x *Populus alba*, clone INRA 717-1B4, was a gift received from Dr. Steve Strauss of Oregon State University. Plants were kept in a growth room using horticultural LED lighting at a constant photosynthetic photon flux density (PPFD) of 250 μmol m^-2^s^-1^, with a 25°C/22°C day night temperature, and humidity levels of 60% relative humidity. Parental plants were shaped to allow each plant to produce a minimum of 10 axillary shoots approximately 10-13 cm in length from which ramets would be collected. A number of cuttings were taken from each parental plant to create sister plants or ramets. Ramets were given a 0.35% IBA treatment then placed in rockwool plugs in trays containing water. Trays were kept in a controlled environment at a constant 24°C temperature, 60% relative humidity, and a PPFD of 100 μmol m^-2^s^-1^ for root initiation. Ramets were given 14 days for root formation.

Once ramets had sufficient root growth they were transplanted into 2 gallon pots and randomly distributed within the growth room. Similar growth conditions to the parental plants were applied to the ramets through the duration of the experiment. The growth room was maintained at a constant photosynthetic photon flux density (PPFD) of 120 or 280 μmol m^-2^s^-1^ measured at a 90 cm height, with a 25°C/22°C day/night temperature, and humidity levels at 60% relative humidity. Ramets were supplemented with 100 ppm nitrogen solution throughout the experiment. Plants were transplanted again after 10 weeks in the 2 gallon pots into 3 gallon pots and moved to a high ceiling growth room for biomass accumulation experiments.

### Agrobacterium-mediated transformation of Poplar hybrid clone 717-1B4

Hybrid *Populus tremula* x *Populus alba*, clone INRA 717-1B4 was a gift received from Dr. Steve Strauss of Oregon State University. Recombinant *Agrobacterium tumefaciens* colonies transformed with pLC0102 were cultured for 24 hours on an orbital shaker. Leaves of the hybrid *Populus tremula* x *Populus alba*, clone INRA 717-1B4 were collected and used to grow stem explants. Stem explants were swirled in the *Agrobacterium* suspensions for 1 hour, and then co-cultured on agar medium for 48 hours at 24°C in darkness. The explants were washed four times in 50ml centrifuge tubes with deionized water at 125 rpm at 24°C, transferred to growth media with antibiotics for 21 days at 24°C in darkness, and then transferred to standard light conditions 16 h light/ 8 h dark to recover green calli. The green calli were transferred to agar medium containing antibiotics for generation and selection of transformed shoots. The regenerated shoots were excised when they were approximately 1 cm long and transferred to agar medium supplemented with antibiotics for rooting. After 5-8 weeks, the rooted and elongated shoots were propagated on the same medium. Rooted LC0102 transgenic plants were transferred out of Magenta boxes in tissue culture into soil pots, acclimated, and grown under an inverted diurnal cycle. Eight weeks post potting, T0 plants went through a vegetative propagation cycle using a stem cutting method frequently used in horticulture practice. Briefly, the apical meristem was removed to release the apical dominance to stimulate the formation of axillary buds for ramet propagation. To track the genetic lineage of transgenic lines, cuttings from the T0 plant are called C1 plants. Cuttings from C1 plants are called C2 plants.

### RNAi design and vector construction

Poplar Plgg1 (Potri.003G099600.1) was identified in *Populus trichocarpa* version 4.1 (Goodstein et al 2012; Tuskan et al 2006) as a homolog to the first plastidal glycolate/glycerate translocator in Arabidopsis (AT1G32080.1) with 73% identity over 96.1% coverage and an E-value of 3.8e-145. To minimize the likelihood of off-target down-regulation, a region of PotriPlgg1 with minimal paralogy was selected. The RNAi region is 256 bp in length, 886-1142 of the transcript and 750-1006 measured from the start codon. RNAi element was designed as an intronic hairpin RNA interference system (ihRNAi) as in the pKANNIBAL plasmid series (Wesley et al 2001), which utilizes the PDK intron between a sense-antisense sequence of the previously defined target region.

Construct LC0102 was designed computationally to resemble the bypass design of South et al. (2019) for expression in poplar using Geneious Prime software (www.geneious.com), synthesized at GenScript (www.genscript.com), and cloned into a binary vector pCambia3200 (Hajdukiewicz et al. 1994) that was then introduced into Agrobacterium GV3101 for poplar transformation.

### Nucleic acid extraction and quantification

100mg of leaf tissues from the second to fourth fully emerged leaf from the apical bud were used for DNA extraction using the NucleoSpin Plant II kit (Macherey-Nagel). Leaf tissues were disrupted with 400µL PL1 from the kit using BeadBeater (BioSpec) at 2400rpm, 1min with four 2.3mm steel beads for 6 times until the tissues are fully homogenized. 10µL of RNase were added to the homogenized tissues then incubated at 65°C for 10mins. Samples were then proceeded with filtration, binding, washing, and elution step based on the manufacturer’s protocol.

30mg of leaf tissues from the eighth fully emerged leaf from the apical bud were used for RNA extraction using the NucleoSpin RNA Plant and Fungi kit (Macherey-Nagel). Leaf samples were sampled from the plants that were at the end of 8 weeks after being transplanted to the 2-gallon pot. Leaf tissues were disrupted with 110µL lysis buffer containing 100µL PFL and 10µL PFR from the kit using BeadBeater (BioSpec) at 2400rpm, 30 secs with four 2.3mm steel beads for 2-3 times until the tissues are fully homogenized. 440µL of lysis buffer containing 400µL PFL and 40µL PFR were added into the homogenized sample then incubated at 56°C for 5mins. Samples were then proceeded with the manufacturer’s protocol with 95mL DNase treatment for 15 mins at RT after PFW1. DNA and RNA concentrations were measured using Synergy LX reader with Take3Trio plate (Agilent) in 2µL volume.

### Primer design and evaluation

Primers and probes were designed using PrimerQuest Tool (Integrated DNA Technologies) with default settings. Gene sequences were obtained from NCBI (https://www.ncbi.nlm.nih.gov/) or AspenDB (http://aspendb.uga.edu/). Each primer set was mapped to the *P. tremula* and *P. alba* sequences obtained from AspenDB using Geneious Prime and manually adjusted to ensure primer specificity and universal amplification from both alleles. The specificity of the primers was further validated using PCR and DNA gel electrophoresis on 1% (w/v) agarose gels containing 1X SYBR Green I Nucleic Acid Gel Stain (Thermo Fisher Scientific) in 1X TAE buffer. Primer efficiencies were evaluated with qPCR or qRT-PCR in singleplex condition using DNA or RNA purified from transgenic plants as standard with 5X dilution series starting from 100ng and 1µg respectively. Standard curves were generated using CFX Maestro software (Bio-Rad) to calculate the primer efficiency. Primer sequence, concentration and amplicon length are listed in Table 1.

**Table 1.**
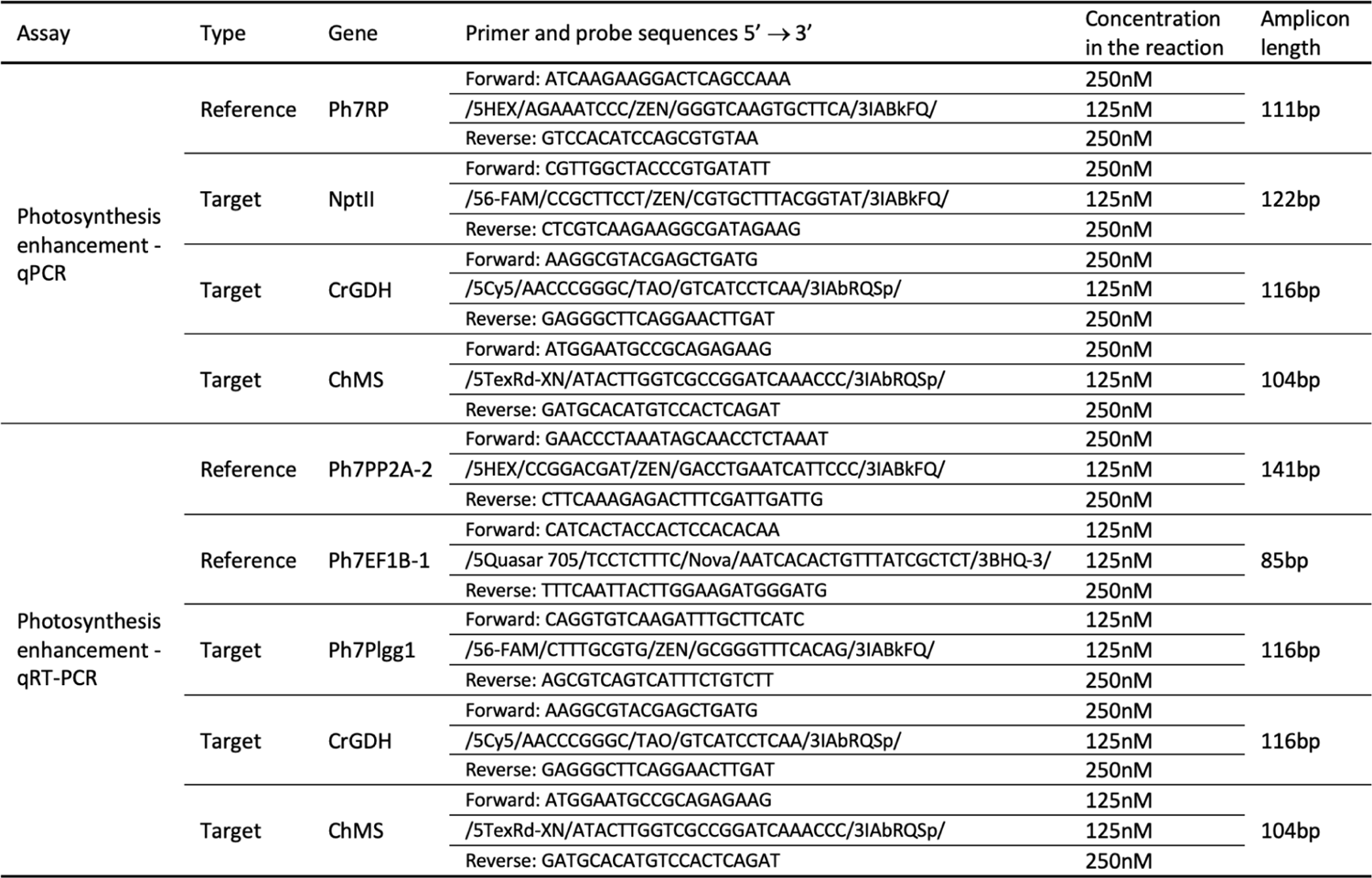
Primer and probe sequences for qPCR and qRT-PCR

### PCR, qPCR, and qRT-PCR

PCR analysis was performed to provide quick validation of transgenic status (Table 2) or primer specificity (Table 1). The 20µL reaction contains 20ng genomic DNA, 1X GoTaq Master Mix (Promega), and 500nM each of forward and reverse primers. PCR was performed with the following conditions: 2 mins 95°C initial denaturation; 35 × (30 secs 95°C denaturation; 30 secs 54.1°C annealing, 2 mins 72°C extension); 5 mins 72°C final extension.

**Table 2.**
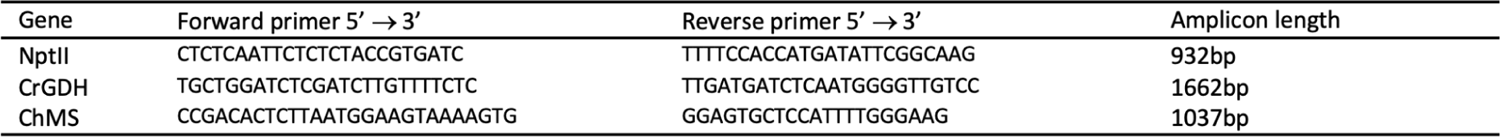
Primer sequences for PCR validation of transgenic status

Multiplex qPCR analysis with technical quadruplicates in each event was used for copy number estimation using CFX Opus 96 machine (Bio-Rad). Multiplex condition has been validated by comparing singleplex and multiplex results to see if the Cq value changes according to the standard mentioned in the primer design section. The 20µL reaction contains 20ng genomic DNA, 1X TaqPath ProAmp Multiplex Master Mix (Thermo Fisher Scientific), primers, and probes (Table 1). qPCR was performed with the following conditions: 10 mins 95°C initial denaturation; 40 × (15 secs 95°C denaturation; 30 secs 60°C annealing/ extension). RP was selected as the reference gene with a copy number of 2. The estimated copy number of the target was calculated using CopyCaller (Thermo Fisher Scientific). Copy number is calculated based on 2^(−ΔΔCq)^×copy number of the target gene in the calibrator, where ΔCq = Cq_target gene_ - Cq_reference gene_, and ΔΔCq = ΔCq_sample_ - ΔCq_calibrator_.

Multiplex qRT-PCR analysis with technical duplicates in each event using CFX Opus 96 machine (Bio-Rad) was used for gene expression analysis, with multiplex condition validated as described previously. The 20µL reaction contains 200ng total RNA, 1X custom made One-Step RT-qPCR Master Mix with lower amount of DTT (Launchworks), primers, and probes (Table 1). qRT-PCR was performed with the following conditions: 10 mins 53°C reverse transcription; 2 mins 95°C initial denaturation; 40 × (15 secs 95°C denaturation; 30 secs 60°C annealing/ extension). Reference genes were selected based on the stability of the selected genes in our target tissue. Ph7Act, Ph7EF1B-1, Ph7EF1B-2, Ph7RP, Ph7PP2A-2, and Ph7UBQ7.2 were tested based on literature (Brunner et al., 2004; Basa et al., 2010; Xu et al., 2011; Pettengill et al., 2012; Wang et al., 2015; Tang et al., 2019). The stability of the reference gene candidates was calculated using CFX Maestro software (Bio-Rad). Average Cq of the selected reference genes were used for ΔCq calculation. The 2^(−ΔΔCq)^ method was used to analyze the expression data.

### Locus number

NptII (MG818373), CrGDH (DQ647436), ChMS (X56948), Ph7Plgg1 (Potri.003G099600), Ph7Act (Potri.001G309500), Ph7EF1B-1 (Potri.001G224700), Ph7EF1B-2 (Potri.009G018600), Ph7RP (Potri.001G342500), Ph7PP2A-2 (Potri.015G068300), and Ph7UBQ7.2 (Potri.005G096700).

### Gas exchange measurements

Photosynthesis activity was measured using a Li-Cor 6800 infrared gas analyser (Li-Cor Biosciences, Lincoln, NE) on poplar plants grown in the controlled environment as described in the plant growth and propagation section. Measurements were performed on various developmental stages of poplar plants depending on the experimental plan. The first mature leaf (generally leaf 8 or 9, counting from the first leaf with a petiole separated from the apical bud) was selected for these measurements and all measurements were taken between 8 am and 2 pm. The leaves were maintained at 25°C, 1000 PAR, with VPD fixed at 1.3 kPa. For photosynthetic CO_2_ response curves or *A-Ci* curves, before starting the measurements leaves were acclimated at 410 ppm CO_2_ for a minimum of 8 minutes, until a steady-state was reached. Once steady-state had been achieved, two data points were logged with a 1.5 minutes interval to record ambient *A_net_* (net CO_2_ assimilation under ambient 410 ppm CO_2_). Following these ambient measurements, the leaf was subjected to a series of CO_2_ concentrations ranging from 0-2000 ppm CO_2_ (410, 400, 300, 200, 100, 50, 0, 400, 600, 800, 1000, 1200, 1500, 2000), and measurements were taken when assimilation reached a steady state rate at each CO_2_ setting. *J_max_* and *Vc_cmax_* were derived from an *A-Ci* curve fitted using the R package “plantecophys” (Duursma, 2015). *J_max_* is the maximum rate of photosynthetic electron transport, *Vc_max_* is the maximum rate of carboxylation), and *A_max_* is the CO_2_ assimilation rate at 2000 ppm CO_2_.

To determine the CO_2_ compensation point, CO_2_ assimilation was recorded at a series of sub-saturating CO_2_ concentrations (150, 120, 90, 70, 50 and 30 ppm) and sub-saturating light levels (250, 185, 120 and 70 PAR). Prior to measurements, leaves were acclimated at 410 ppm CO_2_ and 400 PAR (ambient light levels in the growth room) for a minimum of 8 minutes until a steady state was reached. Throughout the measurements the leaf was maintained at 25°C with a VPD of 1.3 kPa. *A-Ci* curves were fitted for each light level and the CO_2_ compensation point was determined using the Walker-Ort slope-intercept regression protocol with the R package “photosynthesis” (Walker and Ort, 2015).

## RESULTS

### 1. LC102 construct design and transgenic line generation and propagation

Construct LC0102 was designed to resemble the photorespiration bypass pathway described in South et al. (2019) with the following components (Fig. 1): 1) a poplar expression cassette consisting of a gene encoding glycolate dehydrogenase from *Chlamydomonas reinhardtii* (CrGDH, XP_001695381.1) under the control of the *Arabidopsis thaliana* Actin 2 (“Act2”) promoter and terminator (AtAct2, U41998.1), and the *Arabidopsis thaliana* Rubisco Small Subunit (AtRbcS) signal peptide (AY097366.1); 2) a poplar expression cassette consisting of a gene encoding malate synthase from Cucurbita hybrid species *Cucurbita* sp. *Kurokawa Amakuri Nankin* (ChMS, GenBank Accession No. X56948.1) operably linked to the *Zea mays* Suppressor Mutator promoter (Spm_PRO, M25427.1), the AtRbcS signal peptide, and the *Agrobacterium tumefaciens* octopine synthase terminator (OCS_Ter, CP033030.1); 3) a transcriptional unit expressing a small interference RNA design 1 (siRNA1); and 4) a poplar expression cassette consisting of the 35S promoter driven expression of a selectable marker gene, nptII.

**Figure 1.**
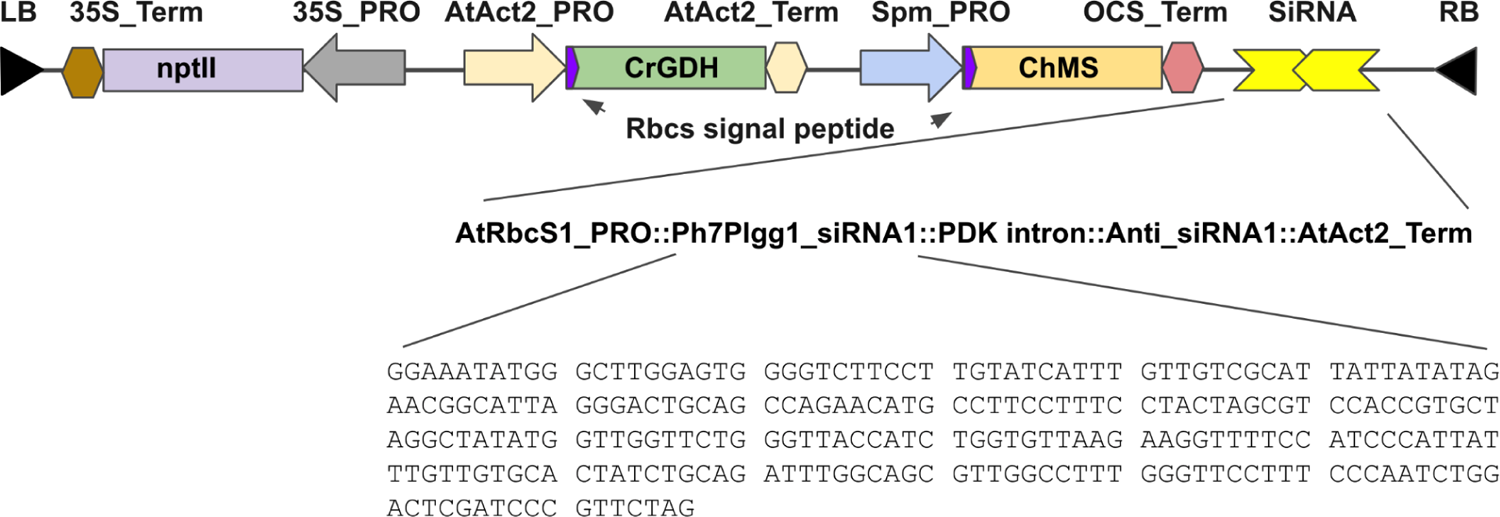
Diagram of construct LC0102 design. Genetic elements within the T-DNA borders are shown. Bottom is the sequence of the sense element of the 256 bp siRNA1 design targeting the region of Ph7Plgg1 transcript from nucleotide position 886 to 1142. An intron of the *Flaveria trinervia* Pyruvate Orthophosphate Dikinase (PDK, GenBank Accession No. X79095.1) was utilized to separate the sense element and the antisense element of siRNA1. Expression of the siRNAi is under the control of the Arabidopsis RbcS1 promoter and the 3’ terminator of the Arabidopsis Act2 gene.

Hybrid poplar INRA 717-1B4 is a hybrid clone of *P. alba* x *P. tremula*. In the diploid genome of the hybrid, any gene would consist of one copy from *P. alba* and one copy from *P. tremula*. Ph7, standing for **P**opulus **h**ybrid **7**17, is used to distinguish the hybrid gene from the parental genes. For example, the Plgg1 gene of INRA 717-1B4 is named as Ph7Plgg1 representing Plgg1 gene sequences from both parental genomes. A database search using the amino acid sequence of Arabidopsis Plgg1 (AT1G32080) as a query against NCBI’s Populus nr database revealed a single hit from *P. alba* with 69.85% AA sequence identity. Using the sequence of PaPlgg1 (Genbank accession# XM_035032358) as query to search for *P. tremula* homologs in the popgenie database, a genome and transcriptome database focused on Populus, resulted in a single hit (Potra000595g04516.1) from *P. tremula* which shares 99.24% AA identity with PaPlgg1. These database searches suggested a single Plgg1 protein in Populus. Comparison of the coding sequences between *P. alba* and *P. tremula* reveals 10 allelic differences with a 6 bp gap and 9 SNPs. The 256 nucleotide siRNA1 design avoided sequences containing the 6 bp gap. To drive the expression of siRNA1, the *Arabidopsis thaliana* Rubisco Small Subunit 1 promoter was selected (AtRbcs21b, X14564.1) (Fig. 1).

We generated 41 independent transgenic events *via* Agrobacterium-mediated transformation of poplar hybrid 717. Rooted transgenic plants were transferred out of Magenta boxes in tissue culture into soil pots, acclimated, and grown in a controlled environment (16h/8h photoperiod, 24/22°C day/night temperature, 60% relative humidity, 120µE). These plants are recognized as the T0 generation following conventional nomenclature. One week post-potting, phenotype evaluation was carried out on a weekly basis. This evaluation included height measurement, root collar diameter (RCD) measurement, leaf number counting, SPAD reading, and morphology recording. Eight weeks post-potting, T0 plants would go through a vegetative propagation cycle to produce clonal cuttings (also known as ramets) from the same mother plant. Under an in-house optimized protocol, these cuttings would normally establish root systems after approximately two weeks and then be transferred to soil pots. Similarly, growth evaluations were performed on these plants, now called C1 (cutting generation 1) ramets, with three or more biological replicates per event.

Leaf tissue samples were collected when T0 plants were transferred out of the Magenta box. Genomic DNA and total RNA were extracted using NucleoSpin Plant II kit and NucleoSpin RNA Plant and Fungi kit respectively (Macherey-Nagel). PCR analysis was performed to provide quick validation of transgenic status of T0 events. Among the 41 T0 events, 3 were identified from PCR analysis as escapes from transformation since they did not contain any of the four transgenes. Gene expression analysis using a SYBR Green qRT-PCR assay showed transgene expression of CrGDH, ChMS, and nptII in the 38 true transgenic events, at various levels. The effect of RNAi design was also evaluated using SYBR Green qRT-PCR assay. Apparent reduction of the target gene Ph7Plgg1 was observed in some of the 38 T0 plants. Some T0 lines showed undesired growth phenotypes such as dwarfing, possibly due to the random insertion of transgenes in the genome that disrupted essential genes. These events were not carried forward for C1 evaluations.

### 2. Growth evaluation of LC0102 transgenic plants in controlled environments

Growth evaluation was performed with 106 C1 plants vegetatively propagated from 30 T0 transgenic events, along with 20 non-transgenic control plants, in a controlled environment (16h/8h photoperiod, 24/22°C day/night temperature, 60% relative humidity, 120µE). Vegetatively produced plants that derived from the same T0 event are clonal variants called ramets. In this C1 growth experiment, each event was represented with 2-4 ramets.

To examine the physiological state of plants after transplanting and estimate the time plants need to fully acclimate, we chose to measure the value of *Vc_max_* of two sets of non-transgenic plants, one consisting of plantlets from tissue culture propagation and the other from vegetative propagation through cuttings. *Vc_max_* is the maximum rate of carboxylation activity of the photosynthetic enzyme RuBisCo. As shown in Fig. 2, photosynthesis activity in hybrid poplar 717-1B4 plants was high during the early phase of growth with *Vc_max_* as high as 82 (µmol CO_2_ m^-2^*S^-1^) in week 2 post transplant, representing a burst of growth most likely due to the nutrient availability in soil. The value of *Vc_max_* gradually decreased during the period of acclimation. After week 6 post transplant, the value of *Vc_max_* fluctuated between 42 to 64. We reasoned that this was an indication that plants had completed the adjustment to the soil growth environment and that the acclimation process was completed, and that the difference in growth rate afterwards was therefore most likely due to genetic makeup rather than differences in transplanting adjustment.

**Figure 2.**
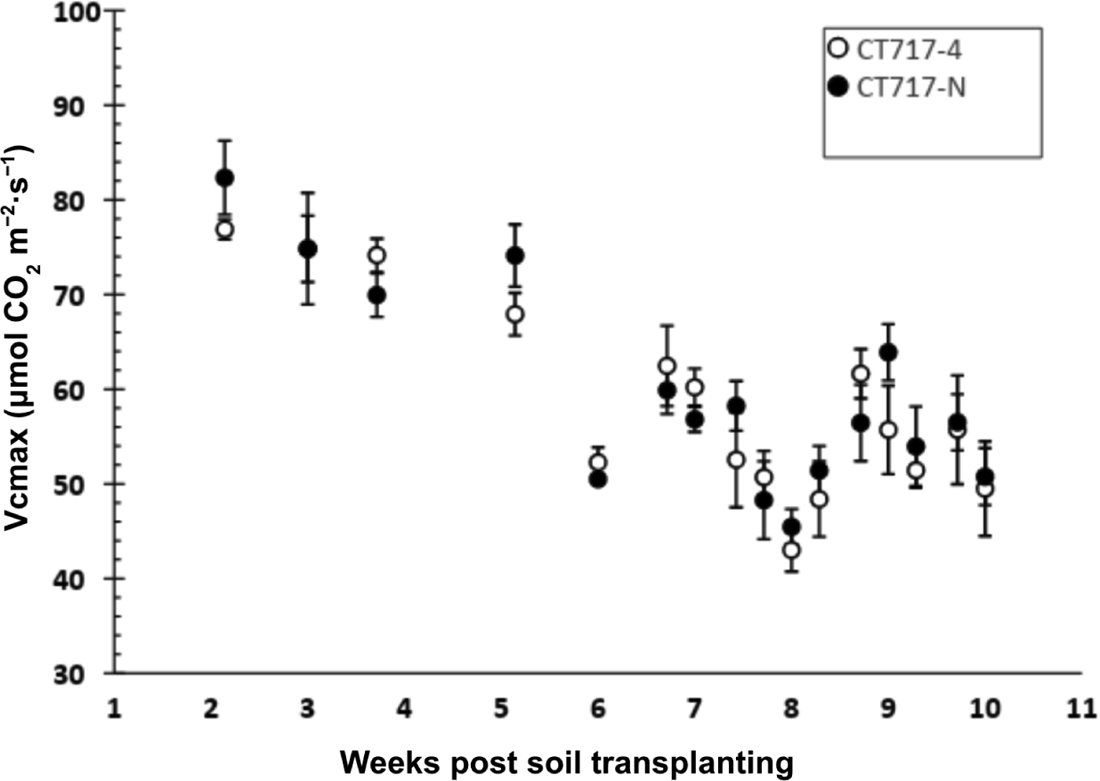
Photosynthesis activity in the form of *Vc_max_* in poplar plants after transplanting to soil pots. Post transplanting, light intensity was gradually increased from 50µE at crown height to 300µE at 50µE per day. Growth conditions: RH, 60%, 16h/8h light/dark cycle, with temperature at 25°C during the day and 22°C at night. Gas-change measurement was performed weekly starting from 2 weeks post soil. More frequent measurement was performed after 6 week post transplanting. *Vc_max_* values were derived from the A-Ci curves. CT717-4: 10 ramets propagated from a soil grown CT717 mother. CT717-N: 10 ramets propagated in tissue culture and directly transferred to soil.

We therefore evaluated the growth rate of 30 T0 events based on height increase over the initial 9 weeks of post planting growth. Fig. 3 shows the height increase and growth rate comparison of a group of 7 T0 events that were transplanted on the same day. At week 9, certain transgenic events such as 7F started to differentiate from non-transgenic control plants (Fig. 3A). A higher rate of growth (cm/week) was also observed in 7F (Fig. 3B). Some transgenic events such as 1A showed disadvantageous growth as indicated by reduced plant height and slowed growth rate. Such cases of negative impact on growth and development in some transgenic events are typical in the process of plant transformation, possibly due to the random insertion position in the genome of an event disrupting some gene functions. Based on the growth evaluation of plant height increase at week 9, a total of 13 transgenic events were selected for advancement to seedling preparation for field planting (additional information in section 7). Transgenic events 13-15E, 5A, 7F, and 8-9A along with non-transgenic control CT717 were carried forward in a high ceiling growth room for further growth evaluation and biomass analysis.

**Figure 3.**
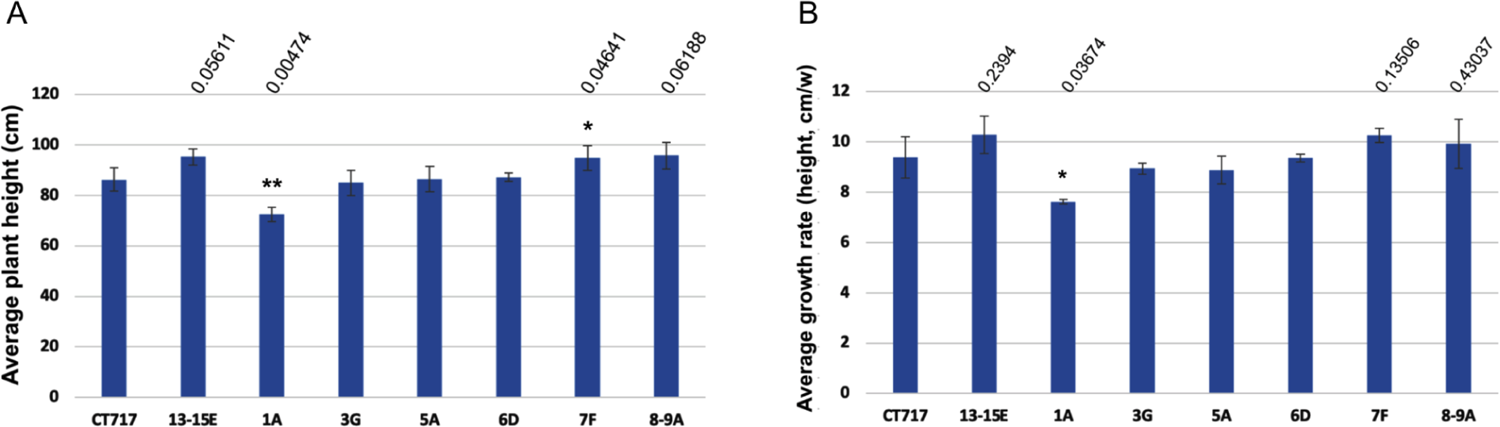
Comparison of plant growth between transgenic events and non-transgenic control 9 weeks post transplanting. Number of ramets: CT717, 5; 13-15E, 2; All others, 3 per event. **A.** Average plant height at week 9. **B.** Average plant height growth rate from week 1 post transplanting (cm/week).

## 3. Molecular evidence for reduction of glycolate transporter expression *via* RNAi technology

### 3.1 The expression level of Ph7Plgg1 is reduced in transgenic plants selected from C1 group

To analyze the effectiveness of our RNAi design on reducing expression of the endogenous glycolate transporter gene Ph7Plgg1 in transgenic events 13-15E, 5A, 7F, and 8-9A, we first developed a multiplex qRT-PCR analysis protocol using a probe assay. Based on literature (Pettengill et al., 2012), Ph7Act was initially chosen as the reference gene for multiplex qRT-PCR analysis. We then analyzed gene expression levels in plants grown in the high ceiling growth room. Fig. 4 shows normalized relative expression level of Ph7Plgg1 in tissue samples collected from the first and the third fully emerged leaf from the apical bud, of these plants. A reduced expression level of Ph7Plgg1 was observed in all four transgenic events indicating that the RNAi design in construct LC0102 functions properly. The degree of reduction in Ph7Plgg1 expression varies from event to event, representing the variation in the expression level of transgenic RNAi among independent transgenic events. In events 13-15E, 5A, and 7F, the level of Ph7Plgg1 reduction is statistically significant in leaf #3 with *p < 0.01* for event 5A and p *< 0.05* for events 13-15E and 7F. This result provides the molecular demonstration that the expression of endogenous gene Ph7Plgg1 in the four transgenic events is inhibited, and suggests that this inhibition along with the expression of the engineered shunt pathway may lead to increased photosynthesis activity and biomass production.

**Figure 4.**
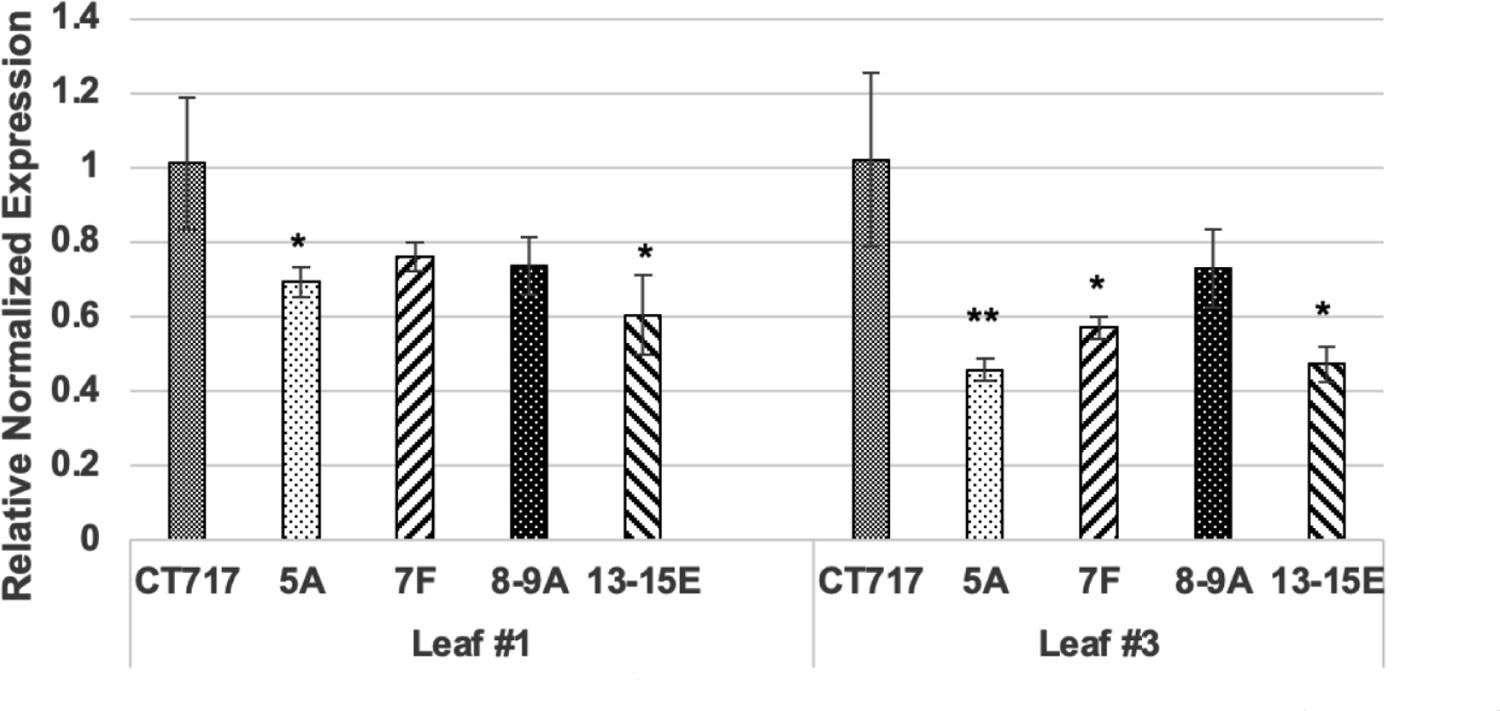
The expression level of Ph7Plgg1 is reduced in transgenic events. Tissue samples were collected from three ramets per event except for 2 for event 13-15E. Reference gene used in this multiplex assay: PtAct (Potri.001G309500). Leaf #1: The first fully emerged leaf; Leaf #3: The third leaf from the growing bud. * and **: Statistically significant at *P* < 0.05 and *P* < 0.01, respectively, when compared to non-transgenic CT717.3. Statistical analysis was performed based on Dunnett’s multiple comparisons test.

### 3.2 Developmental variation in the expression level of endogenous Ph7Plgg1 gene

The level of Ph7Plgg1 expression reduction in transgenic plants is more pronounced in leaf #3 than in leaf #1 which may indicate a higher level of Ph7Plgg1 expression in leaf #3. To understand the developmental and physiological stage where the expression level of our target genes become relatively stable, we collected tissue samples from various leaf positions of poplar plants and analyzed the level of endogenous Ph7Plgg1 expression along with the reference gene PtAct (Fig. 5A). We found the expression level of Ph7Plgg1 is higher in older leaves while the reference gene PtAct is lower. At leaf position 5, the expression level of both Ph7Plgg1 and PtAct become more stable (Fig. 5A), providing the ideal tissue to analyze the efficacy of the Ph7Plgg1 RNAi design.

**Figure 5.**
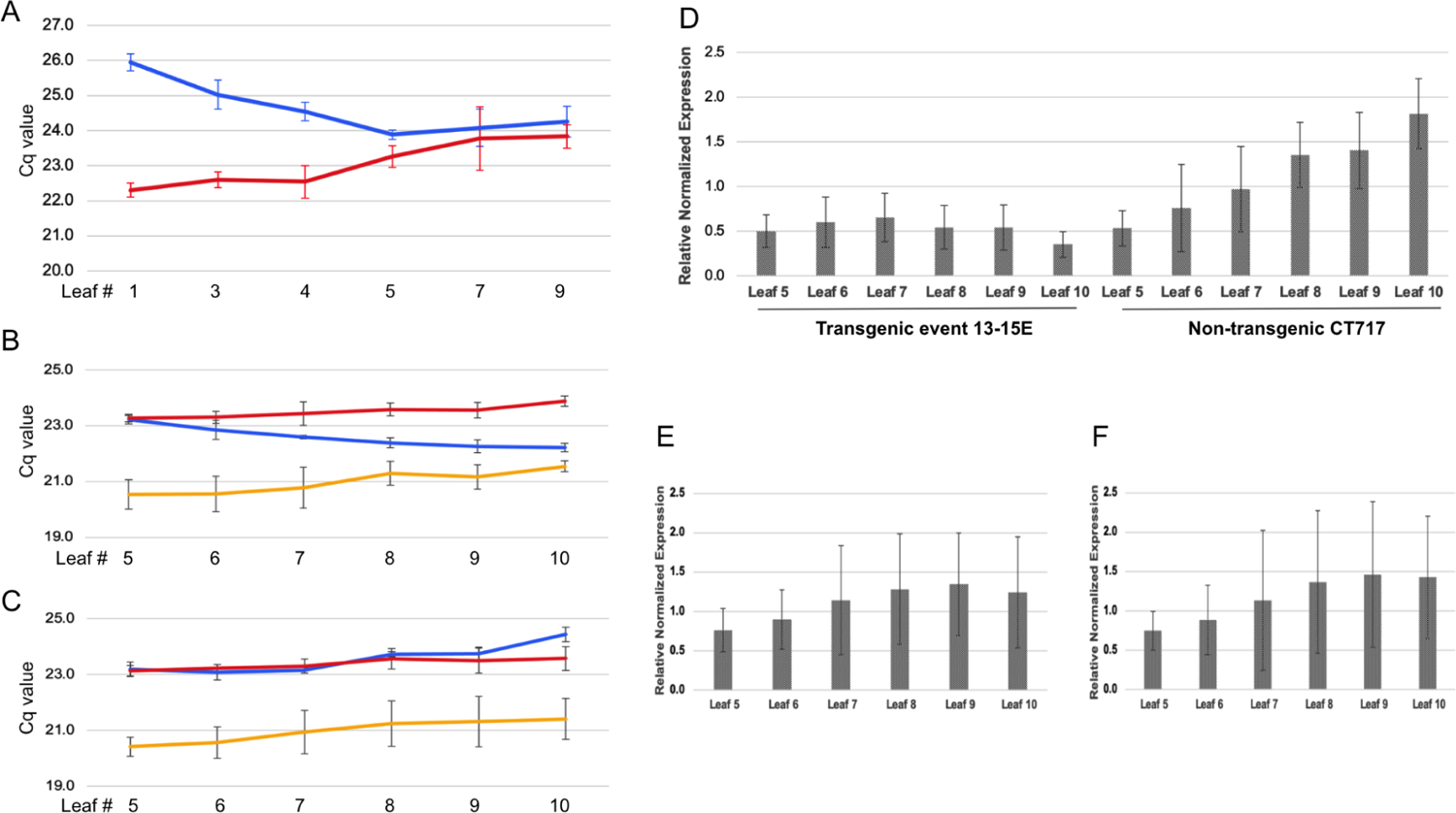
Gene expression variation in leaves of various developmental stages. Total RNA are extracted from 8 weeks old plants grown in a controlled environment as described previously with light intensity at 120µE (**A**) and 280µE (**B-F**). Tissue samples were collected from leaves at each position of three ramets and processed separately to provide three biological replicates for RNA extraction and qRT-PCR analysis. Two technical replicas were employed for the assay. For each reaction, 100ng of total RNA was used. **A.** Absolute Cq value of samples from leaves of various positions on non-transgenic plant CT717. Blue: Ph7Plgg1. Red: Reference gene PtAct. **B & C.** Absolute Cq values of Ph7Plgg1(blue) and two reference genes Ph7EF1B-1 (red) and Ph7RP (yellow) in various leaf positions of non-transgenic plant CT717.3 (**B**) and transgenic plants of Event 13-15E (**C**) grown under 280µE for 8 weeks. **D**. The normalized relative expression level of Ph7Plgg1 in various leaf positions of transgenic event 13-15E and non-transgenic CT717, with the average expression levels of all leaves set as 1.0. **E & F**. Normalized relative expression level of transgenes CrGDH (**E**) and ChMS (**F**) in transgenic event 13-15E. The average expression levels of all leaves was set as 1.0.

A repeat experiment with an increased number of biological replicates was performed to further analyze growth performance (see section 6 for more information) and the level of gene expression. Plants included in the second experiment were grown under an improved growth condition with increased light intensity, 280µE as compared to 120µE for C1 experiment. To search for additional and potentially better reference genes for even more accurate assessment of gene expression in poplar plants, we have conducted database analysis using publicly available RNAseq information and selected 6 candidate reference genes. Gene expression stability and primer efficiency analysis of these genes and their respective probe primer sets have led to two reference genes of choice, Ph7EF1B-1 and Ph7RP, that are more consistent in expression level across various leaf positions (Fig. 5B & 5C). Using these reference genes, further analysis on expression variation of Ph7Plgg1 in leaf positions from 5 to 10 was undertaken. Fig. 5B showed that while the Cq value or relative expression level of the two reference genes are relatively consistent from leaf 5 to 10, the Cq value of Ph7Plgg1 in non-transgenic poplar plants increased from leaf 5 to leaf 10. The Cq value of Ph7Plgg1 in transgenic plants of event 13-15E, however, is relatively more consistent across leaf 5 to leaf 10 (Fig. 5C). Relative normalized expression analysis confirmed the observation from the absolute Cq reading (Fig. 5D). In this analysis, the average expression level of endogenous Ph7Plgg1 in all leaves of non-transgenic control plants was designated as 100% or 1.0 fold. Compared to the average, the expression level of Ph7Plgg1 in other leaf positions varied from 0.53 fold in leaf #5 to 1.82 fold in leaf #10 (Fig. 5D). Relative expression level of Ph7Plgg1 in transgenic event 13-15E is normalized to the average expression of Ph7Plgg1 in all of the non-transgenic control leaves in this test. The leaf position variation in Ph7Plgg1 expression will result in variations in the results of analyses of the effect of RNAi inhibition as shown in Fig. 5D. Variations in the expression levels of the transgenes CrGDH and ChMS were also observed in leaf positions from 5 to 10, although the level of variation was not as high as that of Ph7Plgg1 (Fig. 5E & 5F). Leaf number 8 was selected for tissue sampling to analyze transgene expression and RNAi inhibition.

### 3.3 Reduction of Ph7Plgg1 expression is observed in transgenic plants of the C2 group

Transgenic events 13-15E, 5C, and 7 along with the transformation escape event 16-20 and non-transgenic control CT717.3 were selected for the second experiment that increased the number of ramets per event. Stem cuttings for this experiment were prepared from C1 mother plants and thus were also called C2 plants. Using total RNA extracted from leaf #8 of 8-weeks old plants grown under increased light intensity at 280µE, multiplex qRT-PCR was performed with reference genes Ph7EF1B-1 and Ph7PP2A-2. We found Ph7PP2A-2 performs better than Ph7RP. Normalized relative expression analysis was performed again using 2^(−ΔΔCq)^ methodology. Fig. 6 summarizes the relative expression level of CrGDH, ChMS, and Ph7Plgg1. On average, the relative expression level of transgene CrGDH (Fig. 6A) and ChMS (Fig. 6B) is not significantly different between the three transgenic events 13-15E, 5C, and 7, each with a minimum of 6-9 ramets. For the RNAi target gene Ph7Plgg1, however, there is significant difference between transgenic events and non-transgenic plants (Fig. 6C). The relative expression level of Ph7Plgg1 in non-transgenic poplar was designated at 1.0 fold. Plants of 16-20, a transformation escape that does not contain any transgene, have a relative expression level of Ph7Plgg1 at 1.02 fold, nearly identical to that of non-transgenic CT717.3 plants. The average relative expression level of Ph7Plgg1 in transgenic events 13-15E and 5C, each at 0.24 and 0.35, respectively, is much lower than that of the transformation escape 16-20 or the nontransgenic CT717-3. A greater than 65% reduction in Ph7Plgg1 expression occurs in plants of these two transgenic events. Plants of transgenic event 7 also showed a reduction of Ph7Plgg1 expression, but not at a level that the Tukey’s range test would indicate as significant (Fig. 6C).

**Figure 6.**
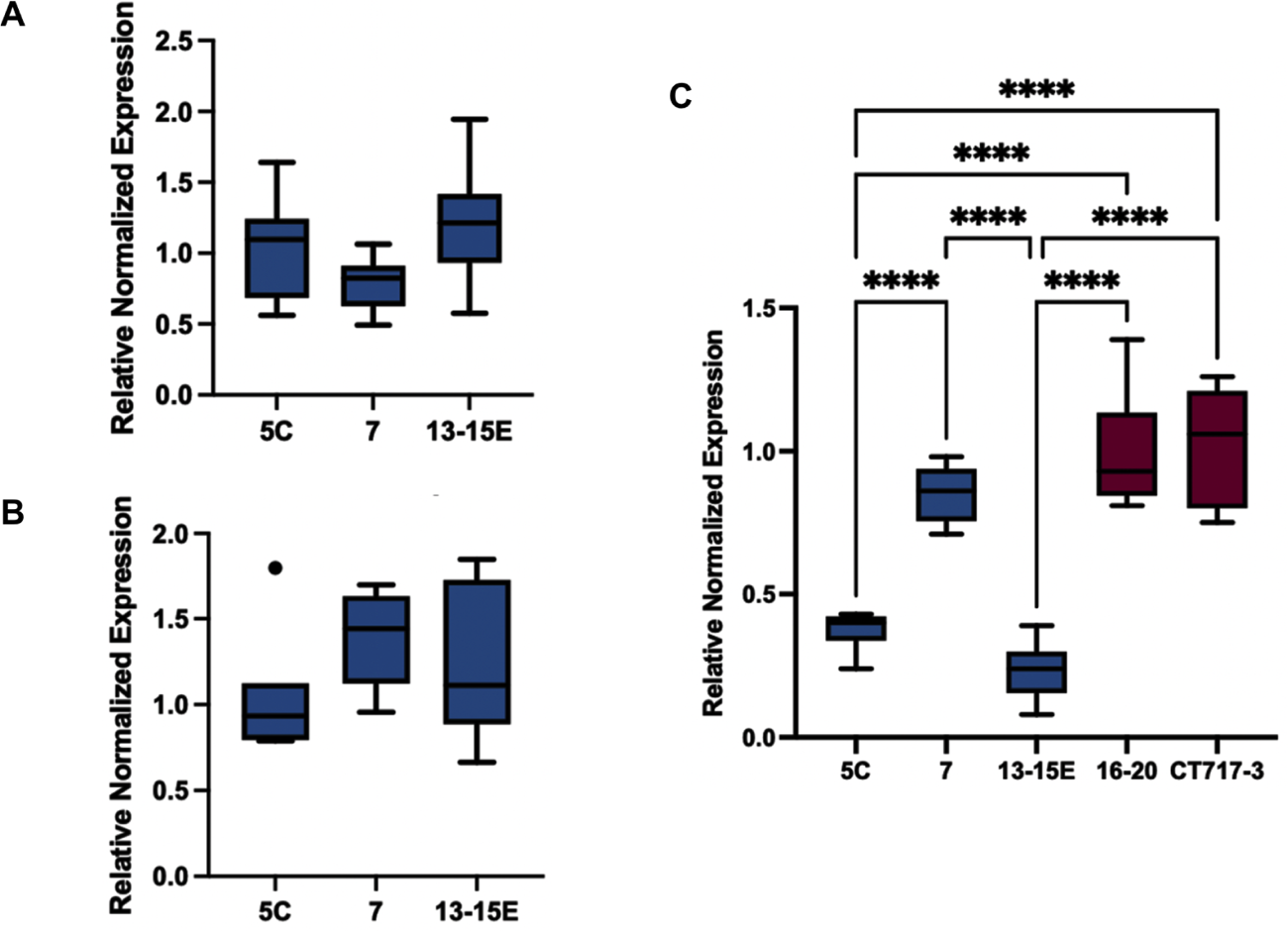
The expression level of Ph7Plgg1 is reduced in transgenic events of the repeat experiment. Tissue samples were collected from leaf 8 of plants 8 weeks post transplanting. Reference genes: Ph7EF1B-1 and Ph7PP2A-2. Number of ramets: CT717-3, 9; 13-15E, 9; 16-20, 9; 5C, 6; Event 7, 8. Technical replicate: 2. ****: Statistically significant between two plant groups at *P* < 0.01 in a pairwise comparison. Statistical analysis was performed based on Tukey’s range test.

In summary, the results of gene expression analysis indicate a significant reduction in the expression level of Ph7Plgg1 in transgenic events 13-15E, 5A, 5C, 7, and 7F. Reduction of Ph7Plgg1 expression would result in lower amounts of glycolate being transported out of chloroplast, a key strategy to inhibit photorespiration. The results also indicate successful expression of CrGDH and ChMS, a shunt pathway to metabolize the retained glycolate back to CO_2_ in the chloroplast for carbon fixation *via* the Calvin cycle. Taken together, these findings led us to anticipate that these transgenic plants would display enhanced photosynthesis efficiency leading to increased biomass production.

## 4. Four-month-old plants of LC0102 events displayed increased growth rate in plant height, stem volume, and photosynthetic rate

### 4.1 Increased stem volume growth rate in transgenic events 13-15E and 7F

After being transferred into large containers (3 gallon) at week 10, C1 plants of transgenic events 13-15E, 5A, 7F, and 8-9A along with non-transgenic CT717 were placed in the high ceiling growth room for further growth analysis. Height and root collar diameter (RCD) measurements continued each week. When trees were tall enough, DBH (diameter at breast high) was measured. Stem volume (V = RC Area × height 1/3) was calculated to estimate differences of biomass accumulation in a young plant. Figure 7 plots the stem volume of transgenic events 13-15E and 7F, along with that of the non-transgenic CT717. As plants grew older, the growth advantage of 7F and 13-15E plants became more apparent compared to non-transgenic plants. Plants of 13-15E started to gain more stem volume and outgrew the 7F plants at week 12 (Fig. 7).

**Figure 7.**
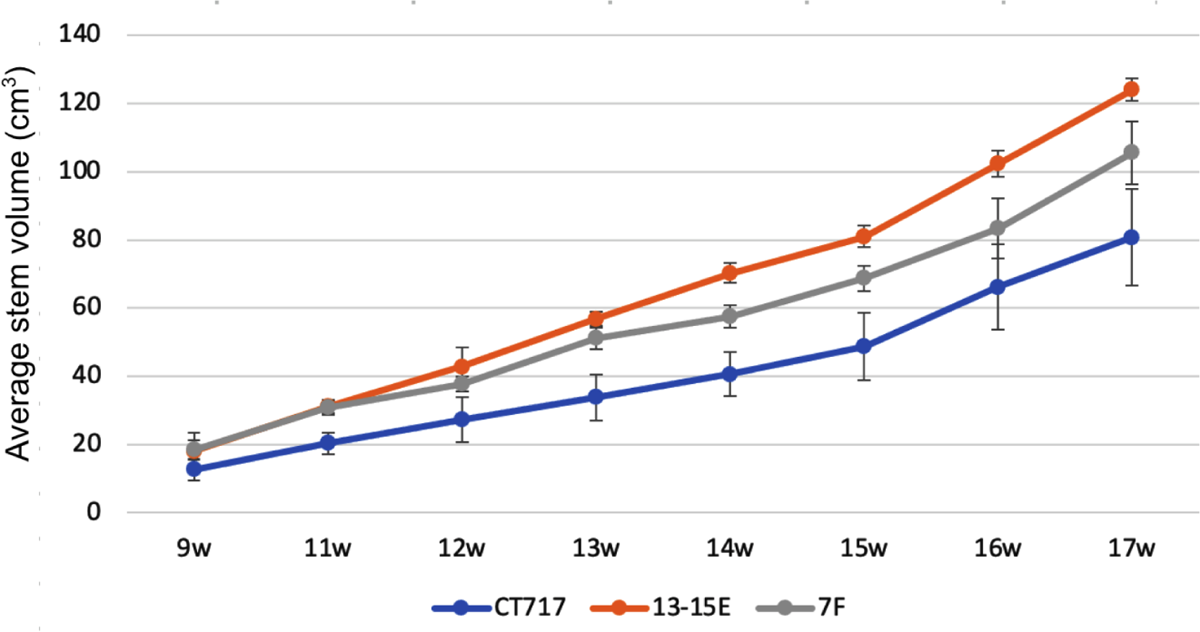
Weekly stem volume growth curves of lead transgenic events 13-15E and 7F plotted against non-transgenic control CT717 from week 9 to 17. Number of ramets: 13-15E, 2; 7F, 3; CT717, 5.

Fig. 8 summarizes the results of growth measurements at week 17, reflecting four months of growth from week 1 when the first set of height data was collected. Compared to non-transgenic control CT717 plants, transgenic plants of event 13-15E and 7F are obviously taller (Fig. 8A) with 18.4% and 14.6% increase, respectively. These plants also deposited more carbon in the stem with increased RCD resulting in significantly higher stem volume as shown in Fig. 8B. The respective stem volume increases over non-transgenic plants at 54% and 31% in events 13-15E and 7F are significantly higher than the percentage increase in sheer height. Interestingly, plants of event 8-9A displayed a significant 10.4% height increase over non-transgenic control plants with *p*-values less than 0.05 (*p = 0.02714*), but the increase in stem volume is not significant. Event 5A showed wide variation in plant height and stem volume with one ramet is as tall as plants of 13-15E. The variation among the three ramets of event 5A is too great to render a lower *p-*value.

**Figure 8.**
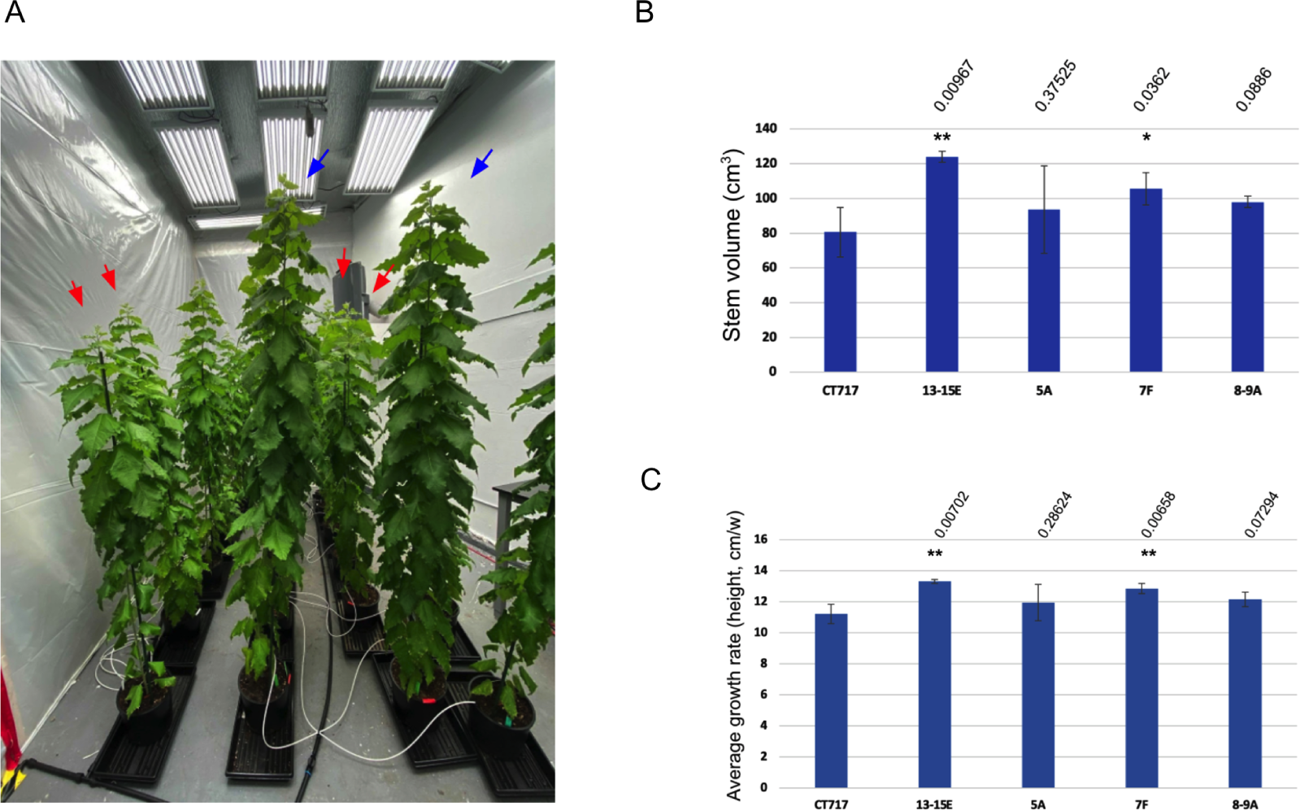
Plant growth analysis of four-month-old C1 plants of events 13-15E, 5A, 7F, and 8-9A in comparison to non-transgenic CT717. Number of ramets: Non-transgenic CT717, 5; 13-15E, 2; 5A, 3; 7F, 3; 8-9A, 3. A. Obvious height difference between transgenic plants and CT717. Blue arrow: 13-15E; Red arrow: CT717. B. Average stem volume of plants at week 17. C. Average growth rate (cm per week) on plant height. * and ** represents a significant difference at *P* < 0.05 and *P* < 0.01, respectively when compared to non-transgenic CT717. Statistical analysis was performed based on one-way ANOVA.

As suggested in Fig. 7, the higher slope in the stem volume growth curve indicates a higher stem volume growth rate in event 13-15E and 7F. The height growth rate increased in transgenic events as well. As shown in Fig. 8C, height increase rates of transgenic events 13-15E, 5A, 7F, and 8-9A are at 13.3, 11.94, 12.84, 12.14 centimeters per week, respectively, compared to 11.21 (cm/w) in non-transgenic control CT717, representing 18.6%, 6.5%, 14.6%, and 8.2% increase, respectively.

### 4.2 Increased CO_2_ assimilation rate in C1 transgenic plants of events 13-15E and 7F

The activity of photosynthesis is typically measured *via* gas exchange which produces A-Ci curves, plots of photosynthetic CO_2_ assimilation versus CO_2_ inside the leaf (to remove any influence of stomata). Using a Li-6800 portable photosynthesis system (LI-COR Biosciences, Nebraska, USA), we measured A-Ci curves of plants grown in controlled environments at week 18. The results indicate that two transgenic events 13-15E and 7F achieve a higher rate of photosynthesis than controls when not limited by light or carbon dioxide (Fig. 9). The higher rate of photosynthesis reflected the increased growth rates in events 13-15E and 7F. The other two C1 transgenic events 5A and 8-9A showed no significant difference in photosynthesis rates when compared to non-transgenic controls (data not shown). We did not observe significant differences in the value of *Vc_max_* between transgenic plants and control, however.

**Figure 9.**
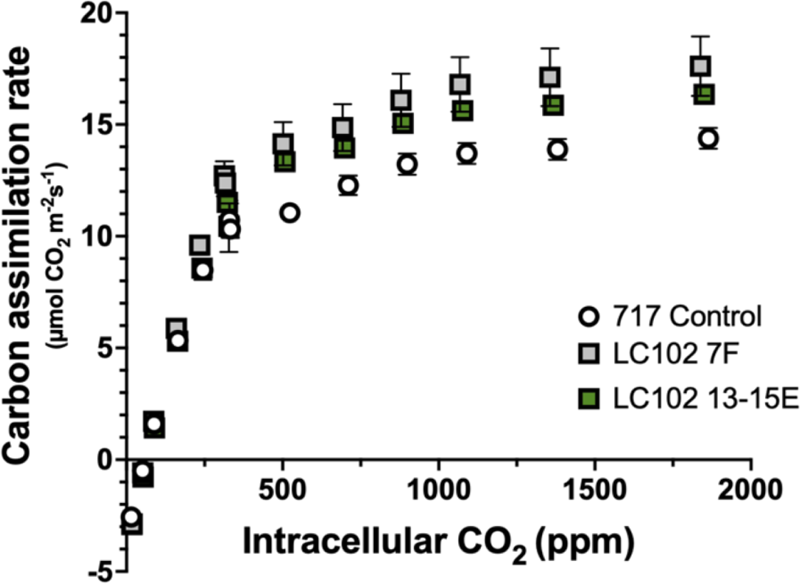
A-Ci curve generated from 18 week old non-transgenic 717 control plants and C1 plants of two transgenic events 13-15E and 7F. Number of ramets measured: 717 control plants, 5; 13-15E, 2; 7F, 3. Error bars indicate SEM.

## 5. Biomass production is increased in C1 plants of transgenic events 13-15E and 7F

The construct design was intended to inhibit photorespiration by reducing the expression of glycolate transporter gene Ph7Plgg1, enabling the metabolism of retained glycolate back into CO_2_ by the expression of CrGDH and ChMS, and collectively increasing the rate of photosynthesis. Our analytical results indicate that Ph7Plgg1 reduction and expression of CrGDH and ChMS was successful (Fig. 4 and 6), the rate of photosynthesis was enhanced in 13-15E and 7F (Fig. 9), and that as a result, the growth rate is increased (Fig 7 and 8). Therefore, it is expected to see increased biomass production in plants of transgenic events 13-15E and 7F in the C1 experimental group.

After 21 weeks of growth in controlled environments, plants of transgenic events 13-15E, 5A, 7F, and 8-9A, along with non-transgenic CT717 plants were harvested for biomass measurements. Leaf, root, and stem of each plant was harvested separately and measured to collect fresh weight (FW). Roots were removed from pots and washed with a gentle message to remove residual soil, after which they were tapped dry on paper towels. After fresh weight measurement, tissues were placed in a pre-weighed and labeled paper bag and placed in a gravity oven to dry at 60°C for 7 days. Each bag of tissues was measured after water and volatile compounds were completely removed. As shown in Fig. 10, C1 plants of transgenic event 13-15E have significantly higher biomass production in all tissue types in both FW and DW, with above ground (AG) DW showing a 53% increase over that of non-transgenic CT717 control plants. AG biomass combines the mass of both leaf tissue and stem tissue. Plants of transgenic event 7F also displayed significantly higher biomass production in stem and roots at both FW and DW levels, but not in leaves. C1 plants of the other two transgenic events, 5A and 8-9A, did not show significantly higher biomass production (data not shown). This is approximately in line with earlier estimation using stem volume (Fig. 8B) or projections using a regression model based on plant height or diameter at breast height.

**Figure 10.**
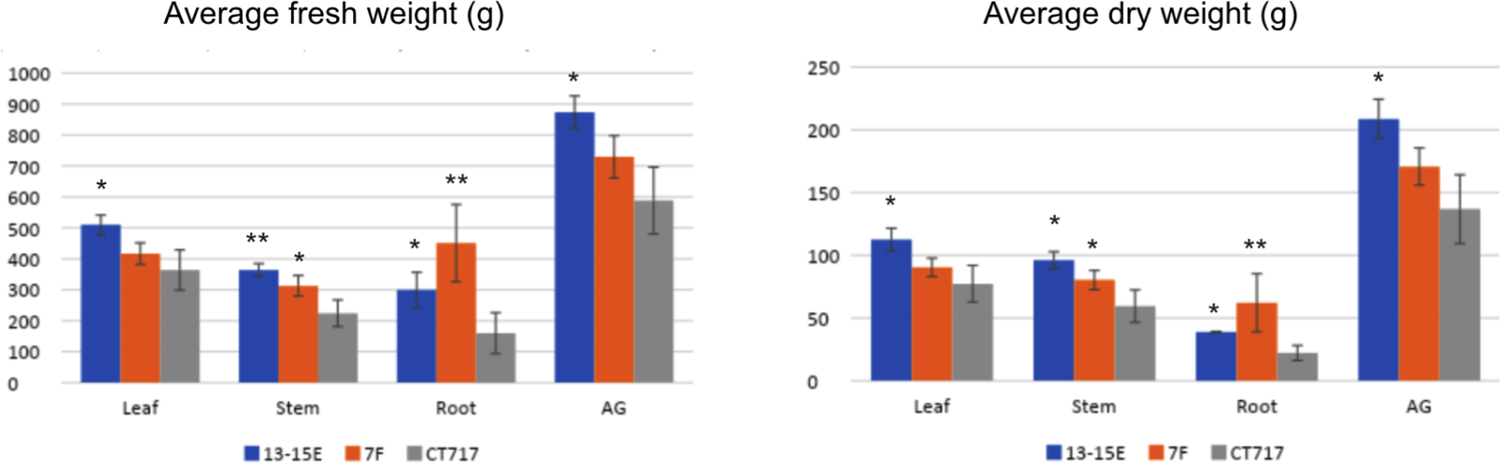
Biomass production increases in C1 plants of transgenic events 13-15E and 7F after 21 weeks of growth in controlled environments. Number of ramets: Event 13-15E, 2; Event 7F, 3; Non-transgenic control CT717, 5. AG: Above ground biomass = Leaf + Stem. * and ** Represents significant difference at P < 0.05 and P < 0.01, respectively when compared with non-transgenic CT717. Statistical analysis was performed based on one-way ANOVA.

## 6. Increases in growth rate and photosynthesis activity observed in transgenic plants of the ongoing C2 repeat experiment

### 6.1 Increased height and stem volume growth in transgenic plants of C2 repeat experiment

To repeat the growth evaluation on transgenic events with an increased number of biological replicates (ramets), a minimum of 7 ramets per event were prepared from C1 mothers of the following events: 13-15E (an event that showed increased biomass production in C1 experiment), 5C and 7 (two transgenic events showing signs of growth advantage at week 9 in C1 experiment), 16-20 (a transformation escape that does not contain transgenes), and the non-transgenic control CT717-3. Plant height and RCD were measured on a weekly basis as in the C1 experiment. Stem volume was calculated using RCD and height as described. Leaf tissue samples were collected for gene expression analysis and as described previously in section 3, expression of transgenes CrGDH and ChMS as well as reduction in the level of Ph7Plgg1 expression were confirmed in transgenic events (Fig. 6).

As observed from the C1 experiment, growth differences between transgenic plants and non-transgenic plants of the C2 repeat experimental group started to appear at week 9 post planting (Fig. 11A). Plants of event 13-15E continued to show increased growth in stem volume compared to plants of non-transgenic CT717-3 or the transformation escape 16-20. Plants of event 7 displayed an even better growth compared to that of 13-15E (Fig. 11A). At week 12 post-transplant, plants of 13-15E and 7 showed significant volume increases over the non-transgenic CT717-3, with *p*-values at 0.008 and 0.01, respectively (Fig. 11B). Plants of event 5C seem to fall into two groups phenotypically and thus it is not surprising to see a wide range on the stem volume of these transgenic plants. The transgenic escape 16-20 showed no difference to the non-transgenic control (Fig. 11B).

**Figure 11.**
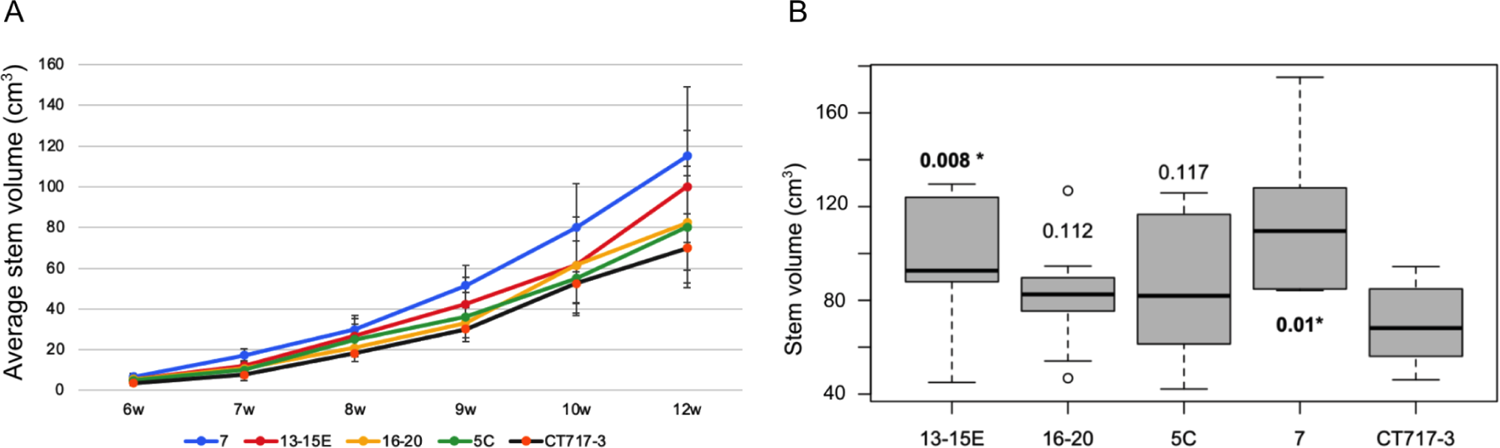
Transgenic events 13-15E and 7 show stem volume increase in the C2 repeat growth experiment. Number of ramets: Event 13-15E, 9; Escape event 16-20, 9; Event 5C, 7; Event 7, 7; Non-transgenic control CT717-3, 9. **A**. Average stem volume over time. **B**. A Box and Whisker plot analysis of stem volume of all 41 plants in the C2 repeat experiment at week 12 post transplanting.

Significant difference in stem volume of 13-15E plants compared to non-transgenic plants exists in the C1 experiment (Fig. 8B). Here we made the same observation again. Along with 13-15E, event 7 also showed higher stem volume growth compared to non-transgenic CT717-3. Plants of event 7 also showed a slightly higher stem volume growth curve compared to that of event 13-15E over the first 12 weeks post transplanting. If the trend continues as it did in the C1 experiment, we anticipate higher biomass production in event 7.

### 6.2. Higher photosynthesis activity in transgenic plants of the C2 repeat experimental group

With the knowledge that poplar plants stabilize from transplanting shock about 8-9 weeks post transplanting (Fig. 2), we started to measure photosynthesis activity of plants in the repeat C2 experiment as early as week 9. *A-Ci* curves were generated from leaf 8 of every plant being measured. *Vc_max_* (the maximum rate of carboxylation), *A_max_* (CO_2_ assimilation rate at 2000 ppm CO_2_), ambient *A_net_* (net CO_2_ assimilation under ambient 410 ppm CO_2_), and *J_max_* (the maximum rate of photosynthetic electron transport) were derived from *A-Ci* curves to provide a broader picture of photosynthesis activity in the measured plant. Less frequently, CO_2_ compensation point was also measured using the Licor 6800 system.

Fig. 12 shows *A-Ci* curves generated from poplar plants of various ages. Previous results indicate poplar plants of 9-11 weeks post transplanting are capable of displaying genetic differences. In the repeat growth experiment, C2 plants of transgenic events 13-15E and 7 showed slightly higher CO_2_ assimilation rate at 9-11 weeks post transplanting (Fig. 12A&B). The difference is small such that the *A-Ci* curves of 13-15E and 7 would overlap each other if they were plotted on the same graph. The difference grew larger when the measurement was performed on plants at 13-14 weeks post transplanting (Fig. 12C). Plants of transformation escape 16-20 showed the same *A-Ci* curve as the non-transgenic control. The *A-Ci* curve of transgenic event 5C shared the same pattern with that of event 13-15E or 7.

**Figure 12.**
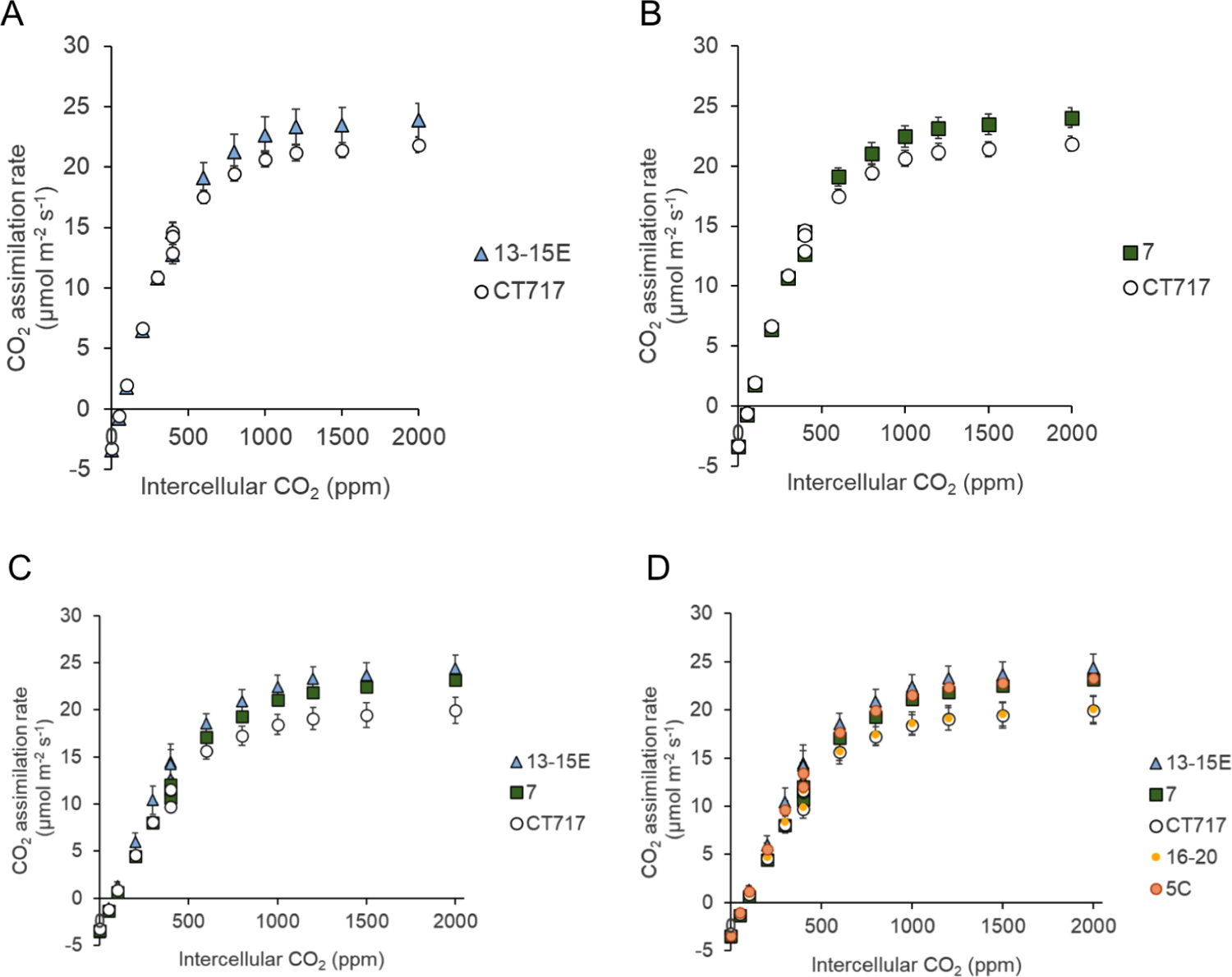
*A-Ci* curves of transgenic plants plotted together with that of non-transgenic plants of the same developmental stage. A & B. 9-11 weeks old transgenic plants of event 13-15E and 7 show slightly higher CO_2_ assimilation rate. Number of ramets: Event 7, 5; Event 13-15E, 7; CT717, 7. C & D. A-Ci curves of 13-14 weeks old plants. Number of ramets per event: 3.

Photosynthetic activity increase in transgenic events 13-15E and 7 is also reflected in *J_max_*, the maximum rate of photosynthetic electron transport for a given light intensity, as shown in Fig. 13A. *Vc_max_* measures the maximum rate of carboxylation activity of RuBisCo. No significant differences in *Vc_max_* of plants in this C2 repeat experimental group were observed. A significant difference was observed in the ratio of *J_max_*:*Vc_max_* between transgenic plants and non-transgenic controls (Fig. 13C). The *J_max_*:*Vc_max_* ratio is an indicator of photosynthesis activity. In transgenic plants of event 13-15E and 7F, the ratio of *J_max_*:*Vc_max_* is significantly higher than that of the non-transgenic control CT717 or the transformation escape 16-20. For transgenic event 5C, even though the *J_max_* and *Vc_max_* are similar to that of the non-transgenic control CT717 or the escape 16-20, the ratio of *J_max_*:*Vc_max_* appears to be higher than both controls with p value at 0.063. As mentioned previously, there appear to be two groups of plants within event 5C morphologically. It remains to be seen if the growth difference between the two groups is due to genetic changes and if that manifests in the ratio of *J_max_*:*Vc_max_*. As shown in Fig. 12, A-Ci curves measured from plants of event 13-15E and event 7 showed higher CO_2_ assimilation rate at higher CO_2_ concentrations. As expected, *A_max_* in 13-15E and 7 is significantly higher than in CT717 or 16-20 (Fig. 13D).

**Figure 13.**
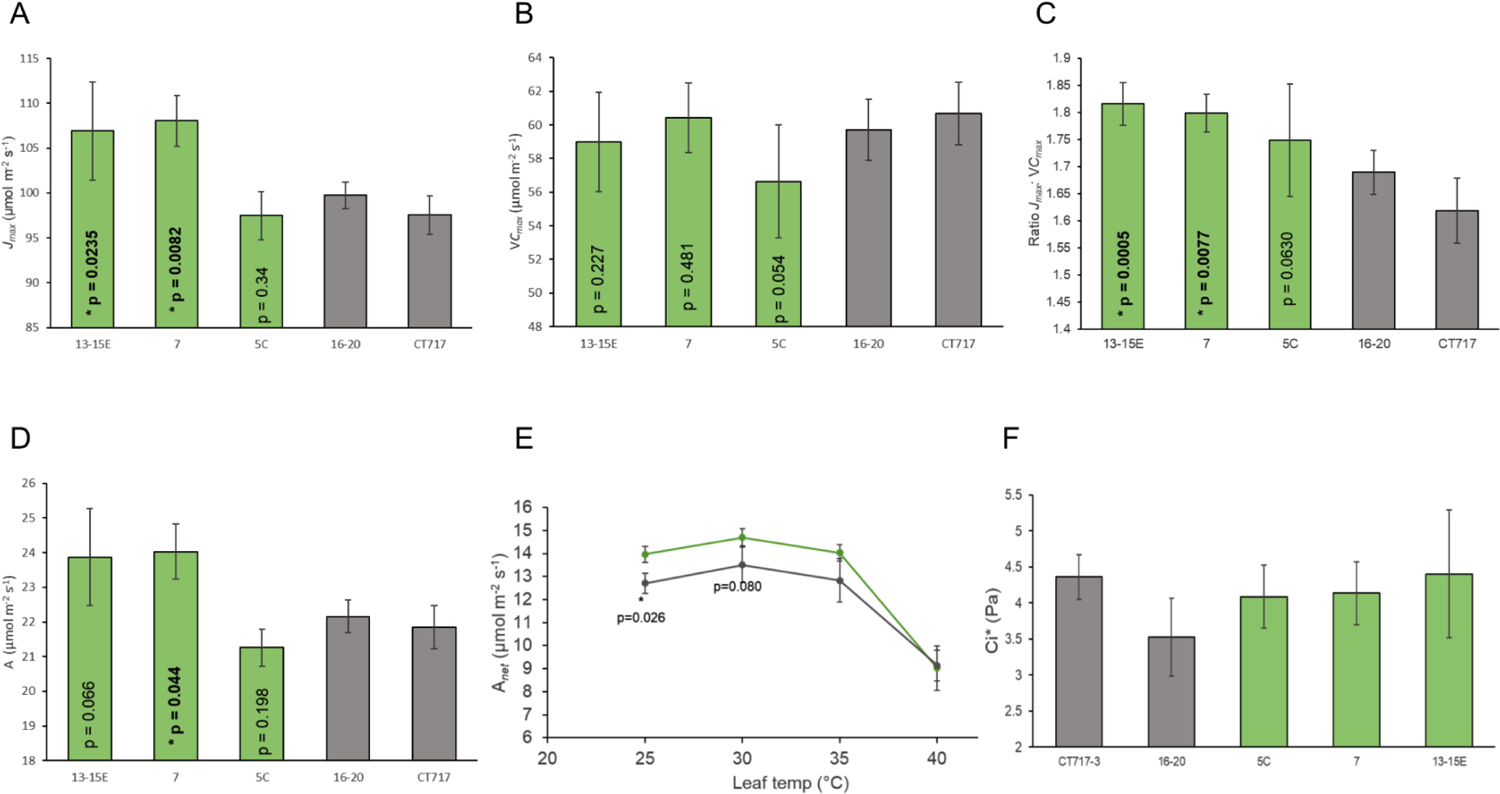
Photosynthetic activity parameters derived from gas exchange measurement. Statistical analysis was performed using a one-tailed t-test, compared to CT717. **A.** *J_max_* of 9-11 weeks old plants. **B.** *Vc_max_* derived from *A-Ci* curve analysis. **C.** *J_max_:Vc_max_* ratio of 9-11 weeks old plants. **D.** Maximum CO_2_ assimilation rate at saturated CO_2_ concentration. **E.** Ambient CO_2_ assimilation increased in 13-15E (green) compared to non-transgenic CT717 (gray) at temperatures as high as 35°C. **F.** CO_2_ compensation point in plants of 12-13 weeks old.

It has been observed that the impact of higher temperature on photorespiratory loss and thus crop productivity loss are mitigated by the less-energy intensive glycolate metabolism pathway similar to our LC0102 design (Cavanagh et al, 2021). We therefore measured the net CO_2_ assimilation rate of 13-15E plants across a range of temperatures from ambient (25°C) to as high as 40°C. *A_net_* at ambient temperature in 13-15E plants is significantly higher than in non-transgenic CT717 plants (Fig. 13E). At increased temperatures of 30°C and 35°C, *A_net_* in 13-15E plants remains higher than that in non-transgenic plants. At 40°C, *A_net_* in both groups of plants decreased dramatically and no difference was observed between the two groups. No significant difference in the CO_2_ compensation point was observed between transgenic and non-transgenic plants (Fig. 13F).

## DISCUSSION

A number of photorespiration bypass pathways have been conceived and tested in C3 annuals (Kebeish et al., 2007; Carvalho, 2011; Maier et al., 2012; Nolke et al., 2014; South et al., 2019; Shen et al., 2019; Wang et al., 2020) thanks to the advancement of knowledge in synthetic biology. Significant increases in photosynthetic efficiency and crop grain yield has been observed (Shen et al., 2019; Wang et al. 2020). After evaluation of design principles, we selected to test a pathway (South et al., 2019) in conjunction with glycolate transporter inhibition in poplar hybrid INRA 717-1B4. Results of molecular analysis and photobiology measurement supported our observation that this bypass increased biomass production in poplar by as much as 53% higher than plants that do not have the functional bypass pathway (Fig 10). To our knowledge, this is the first time that photosynthetic efficiency in a tree species has been enhanced by such a magnitude.

We reasoned that for this photorespiration bypass to work in poplar, reduction in the expression level of the glycolate transporter is critical. As stated before, INRA 717-1B4 is a hybrid clone of *P. alba* x *P. tremula*. Comparison of the coding sequences of Plgg1 between *P. alba* and *P. tremula* reveals 10 allelic differences with a 6 bp gap and 9 SNPs. Our RNAi design avoided sequences containing the 6 bp gap. The various levels of Plgg1 reduction in our transgenic events (Fig. 6C) validated the RNAi design principle. The wide range in reduction of Plgg1 expression in the transgenic events would also allow for selection of events with the appropriate amount of glycolate transportation inhibition. However, plant gene expression is well known to vary with growth and developmental stages as well as physiological changes due to (a)biotic environmental changes. Our qRT-PCR analysis was established and optimized through consideration of factors including the choice of reference genes, the age of plants when samples were collected, and growth conditions-especially light intensity, etc. This analysis protocol enabled minimization of plant-to-plant variations.

In addition to the range of Plgg1 expression reduction, it is desirable to also have a range of expression levels for the two transgenic metabolic enzymes. This allows selection of a balanced perturbation of the metabolic flow that would enable an optimized output from the shunt pathway for increased photosynthetic activity. We observed a range of CrGDH and ChMS expression levels in the 41 T0 events. The selected events for the two growth experiments did not show significant differences in the expression level of CrGDH or ChMS, probably an outcome of growth based selection. Fine-tuned metabolic perturbation is also achieved at the level of post-transcriptional control, such as feedback inhibition and modulation. It would be interesting to quantitatively analyze the levels of related proteins in the transgenic plants. Furthermore, enzymatic activities, governed by enzyme kinetic characteristics, provide another degree of fine-tuning regulation on metabolic output. Enzyme engineering through computational design using software to discern the relationship of protein structure and function, such as Rosetta, has been widely applied in the field of synthetic biology (Leman et al. 2020; Pan and Kortemme, 2021). With the ever increasing data on structure-function relationship and the knowledge from machine learning on enzyme activities, the future of protein design for biotech applications can only improve. Enzyme engineering through directed evolution (Arnold, 2019) to accumulate beneficial amino acid sequence changes, thereby improving enzymatic activity, has proven to be practical and fruitful in synthetic biology applications (Siehl et al. 2014; Piatkevich et al. 2018; Sachsenhauser and Bardwell, 2018). Examples of using DE to improve photosynthetic efficiency have been reported albeit not as successful (Mueller-Cajar and Whitney, 2008). Challenges are not with the technology itself, but rather with how to properly design and screen the enzyme variants that would exert the activity change in a living plant. Additionally, changes in the perturbation of endogenous metabolic flow can be monitored via metabolite profiling experiments or metabolomics analysis. It remains unclear what changes in the metabolite profile of events 13-15E and 7F contribute to the increased biomass production.

When vegetatively propagated plants experience changes in environmental and growth conditions, ranging from mechanical damage, to lack of nutrients during rooting, to transplanting shock, they need a period of time post-potting for acclimation and to stabilize, a term used in the field to describe when genetics start to have the dominant effect on growth characteristics of plants. The growth performance of C1 plants showed a wide range of variation during the first few weeks of growth after transplanting in the growth room. Our results, based on *Vc_max_* measurement of photosynthesis activity in poplar plants after transplanting, helped to evaluate stabilization at the physiological level. Additional research is needed to better understand the process of stabilization and to narrow down the time frame poplar needs to fully acclimate to soil growth.

The impact of engineered photorespiration bypass on the activity of photosynthesis can be measured with conventional gas-exchange methodologies. Changes in *J_max_* and the ratio of *J_max_*:*Vc_max_* reflect changes in photosynthetic activity. For a given plant species, *Vc_max_* fluctuates with the physiological state of the plant and may also change under the influences of growth and environmental conditions, such as (a)biotic stresses, including transplanting acclimation (Fig. 2). However, for plants of the same species grown under the same growth conditions, *Vc_max_* fluctuations should not be significantly different from each other as shown in Fig. 13B.

Cavanagh et al. (2021) showed a greater difference in CO_2_ assimilation rate at higher temperatures for tobacco plants expressing a similar photorespiration bypass pathway. Our engineered poplar plants showed higher CO_2_ assimilation from ambient temperature to 35°C, but no difference at 40°C (Fig. 13E). Because the temperature conditions in our measurement are more transient (15 minute acclimation in the measurement chamber as opposed to 24hrs of acclimation for tobacco plants), the different observations between our engineered poplar and the engineered tobacco of Cavanagh et al. (2021) may not precisely represent the true difference. Alternatively, the inherent difference between the two plant species could also contribute to a different temperature response curve. In addition, we did not detect a significant difference in CO_2_ compensation point between our transgenic poplar and control when plants are 13 weeks post-transplanting. At the time of measurement, due to space constraints, these plants were too close to the light source and may have been under light stress. Since the difference in CO_2_ compensation point can be small (South et al. 2019; Cavanagh et al., 2021), response to light stress might have masked the difference in CO_2_ compensation point.

Through testing of the selected photorespiration bypass pathway in hybrid poplar, we obtained transgenic trees that accumulated up to 53% more above ground biomass during 5 months of growth in a controlled environment. The dry weight of root tissue showed an even greater increase in biomass. However, due to the potential carryover of residual soil, the results from three biological replicates in the C1 experiment may be skewed. Biomass measurement results from the C2 experimental group will help validate the observation on root biomass differences. The next step is to test the growth performance of these plants in field conditions. In order to make the summer field planting in 2021, a total of 13 events were selected from a 9-week preliminary growth evaluation of 30 transgenic events that was performed in the C1 growth experiment. These include 11 transgenic events harboring a varied copy number of transgenes (Fig. 14) and 2 transgenic escapes (event 16-20 and 8-9D) that went through the transformation process but do not contain a transgene. Copy number variation is typical in transgenic events. The field experiment is a collaboration between Living Carbon and Oregon State University (OSU). In mid-July of 2021, 672 poplar seedlings representing the selected 11 transgenic events, 2 transgenic escapes, and non-transgenic control were planted in the OSU field site at Corvallis in two blocks-one large and one small. The large block was planted one week earlier than the small block. The large block has in total 480 seedlings including 372 testing trees and 108 border trees. Despite the high temperature conditions in the summer, there was a very low rate of post-planting mortality (Fig. 15). The photosynthetic performance of these events will be evaluated in the year of 2022.

**Figure. 14.**
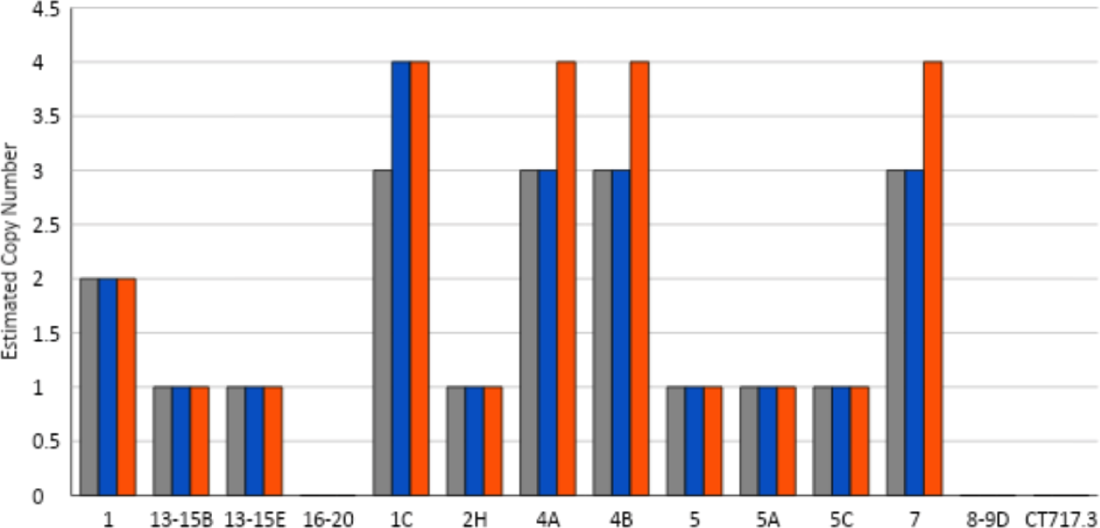
Transgene copy numbers for LC0102 transgenic events included in the field trial. Gray, nptII; Blue, CrGDH; Red; ChMS.

**Figure. 15.**
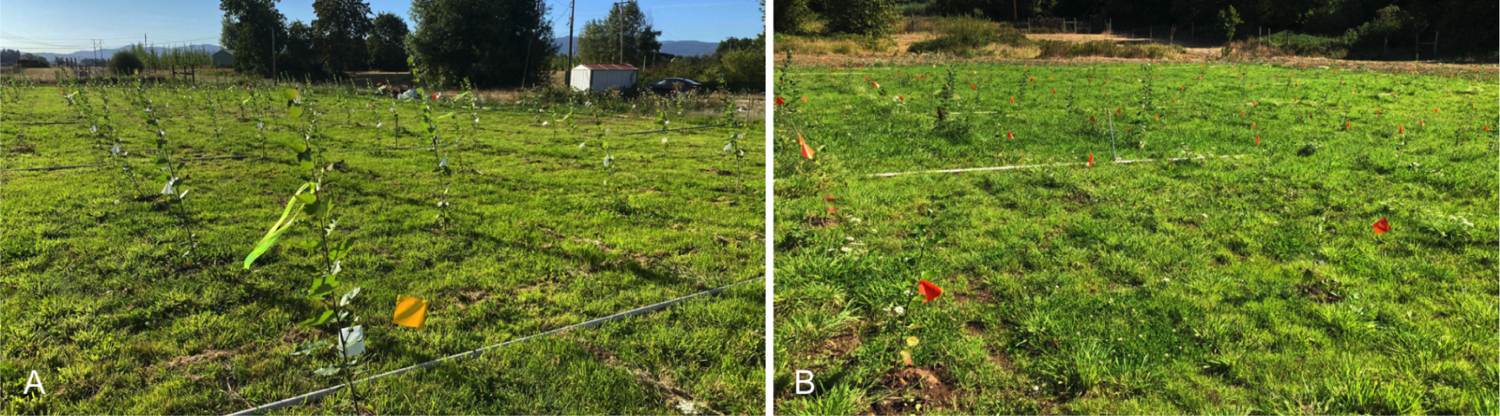
LC0102 trees in the field approximately two months post-planting. A. Large block. B. Small block.

The increase in biomass production in these engineered poplar trees may not fully reflect the efficiency enhancement in photosynthesis. As the amount of photosynthetic carbohydrates increases in the source tissue, transportation and deposition of these energy rich carbon molecules to the sink tissue become limiting factors. Alternative photorespiration bypass pathways have been tested in other C3 plant species. In some cases, the pathway did not work as intended (Carvalho, 2011). In other cases, the pathway seemed to work properly and resulted in enhanced photosynthetic rate, increased biomass and grain yield. However, the seed setting is reduced (Shen et al., 2019; Wang et al., 2020). Through an integrated analysis of transcriptomics, physiology, and biochemistry, Wang et al. (2021) concluded that photosynthetic carbohydrates in these plants were not transported to grains in an efficient manner. In woody plants such as poplar, this source-sink carbon partitioning issue may become limiting when photosynthesis is enhanced. Other issues such as C-N balance may add to the complexity as well. Nonetheless, it may not be as complicated as it is in cereal crops due to complexities in regulating the transition from vegetative growth to reproductive growth for grain production.

As alluded to earlier, we aim to further improve photosynthesis efficiency in poplar, as well as expand application to other woody plant species with the goal to greatly enhance trees’ ability to drawdown atmospheric CO_2_. To make a greater impact on the goal of carbon neutrality and negativity, we are taking a multidisciplinary approach to actively engage in research areas such as decreasing wood decomposition rate to achieve prolonged carbon storage and better yet, permanent carbon storage. The biological system is a powerful one when it comes to carbon drawdown and storage, but at the same time, is complex. Needless to say, it is a challenging goal to engineer trees to make a meaningful impact on climate change. Utilization of the growing knowledge base to test potential strategies in trees is at least a good start.

